# Temporal dynamics and metagenomics of phosphorothioate epigenomes in the human gut microbiome

**DOI:** 10.1101/2024.05.29.596306

**Authors:** Shane R Byrne, Michael S DeMott, Yifeng Yuan, Farzan Ghanegolmohammadi, Stefanie Kaiser, James G. Fox, Eric J. Alm, Peter C Dedon

## Abstract

**Background:** Epigenetic regulation of gene expression and host defense is well established in microbial communities, with dozens of DNA modifications comprising the epigenomes of prokaryotes and bacteriophage. Phosphorothioation (PT) of DNA, in which a chemically-reactive sulfur atom replaces a non-bridging oxygen in the sugar-phosphate backbone, is catalyzed by *dnd* and *ssp* gene families widespread in bacteria and archaea. However, little is known about the role of PTs or other microbial epigenetic modifications in the human microbiome. Here we optimized and applied fecal DNA extraction, mass spectrometric, and metagenomics technologies to characterize the landscape and temporal dynamics of gut microbes possessing PT modifications.

**Results:** Exploiting the nuclease-resistance of PTs, mass spectrometric analysis of limit digests of PT-containing DNA reveals PT dinucleotides as part of genomic consensus sequences, with 16 possible dinucleotide combinations. Analysis of mouse fecal DNA revealed a highly uniform spectrum of 11 PT dinucleotides in all littermates, with PTs estimated to occur in 5-10% of gut microbes. Though at similar levels, PT dinucleotides in fecal DNA from 11 healthy humans possessed signature combinations and levels of individual PTs. Comparison with a widely distributed microbial epigenetic mark, m^6^dA, suggested temporal dynamics consistent with expectations for gut microbial communities based on Taylor’s Power Law. Application of PT-seq for site-specific metagenomic analysis of PT-containing bacteria in one fecal donor revealed the larger consensus sequences for the PT dinucleotides in Bacteroidota, Firmicutes, Actinobacteria, and Proteobacteria, which differed from unbiased metagenomics and suggested that the abundance of PT-containing bacteria did not simply mirror the spectrum of gut bacteria. PT-seq further revealed low abundance PT sites not detected as dinucleotides by mass spectrometry, attesting to the complementarity of the technologies.

**Conclusions:** The results of our studies provide a benchmark for understanding the behavior of an abundant and chemically-reactive epigenetic mark in the human gut microbiome, with implications for inflammatory conditions of the gut.

## Introduction

There are now dozens of enzymatically-installed DNA modifications – the epigenome – in all forms of life, with the greatest diversity occurring in prokaryotes and bacteriophage [1]. While bacterial DNA modifications are best known from restriction-modification (RM) systems, they are now also known to regulate gene expression [2–10]. Phosphorothioation (PT) of DNA, in which a sulfur atom replaces a non-bridging oxygen in the sugar-phosphate backbone, is the only known naturally occurring DNA backbone modification. Developed more than 50 years ago as a synthetic modification to engineer nuclease resistance into oligonucleotides [11], we more recently discovered that PTs occur naturally in the genomes of bacteria [12] and archaea [13]. To date, PTs have only been detected in prokaryotes and only in DNA [14].

Synthesis of PTs is so far known to be mediated by three gene families – DndABCDE, SspBCD, and BrxPCZL (BREX type 4) – whether functioning in RM or regulating gene expression [12, 13, 15–17]. As atypical RM systems, DndFGHI and SspFGH function as restriction endonuclease complexes to cleave unmodified “non-self” DNA [13, 16, 18] in spite of only ∼10-15% of available consensus sequences being modified with PT [19], while variant SspE and BREX type 4 systems possess antiviral activity without restriction endonuclease activity [15, 17]. Similar to methylation-based restriction-modification systems, the Dnd and Ssp modification proteins catalyze phosphorothioation on one or both DNA strands of specific consensus sequences. For example, DndABCDE modifies both DNA strands at 5’-G*AAC-3’/5’-G*TTC-3’ sequences (where “*” denotes a PT linkage) in *Escherichia coli* B7A and 5′-G*GCC-3′/5′-G*GCC-3′ in *Pseudomonas fluorescens* pf0-1 [16, 19, 20], while PTs occur as a single-strand modification at C*CA in *Vibrio cyclitrophicus* FF75 [15, 19].

An unusual feature of PT modification systems is that many PT-containing bacteria lack restriction genes, which suggests an alternative epigenetic role of PTs [19]. For example, PTs have been proposed to provide epigenetic regulation of transcription of redox homeostasis genes [16], which may relate to a signaling function for the easily oxidized and nucleophilic sulfur in PTs. Indeed, there is evidence that PTs provide some protective effects in cells exposed to reactive oxygen and nitrogen species, such as peroxides [16, 21] and peroxynitrite [22]. Contrasting with this protection, PT-containing bacteria are 5-fold more sensitive to neutrophil-derived hypochlorous acid (HOCl) due to extensive DNA breaks at PT sites [23]. Given the widespread distribution of PT gene systems among bacteria and archaea and a preliminary report by Sun *et al*. [24] of PTs present in human fecal DNA, the unusual chemical properties of PTs raise questions about how PT-containing microbes might behave in the healthy gut microbiome or be altered by chronic inflammation of the gut, such as inflammatory bowel disease (IBD) [25, 26].

Here we undertook a foundational analysis of PT epigenetics in the human gut microbiome by exploring the landscape and temporal dynamics of PT dinucleotides in fecal DNA from 11 healthy humans, using an enhanced fecal DNA extraction method and optimized chromatography-coupled mass spectrometry (LC-MS). Data for PTs were compared to a widespread and well established microbial epigenetic mark, m^6^dA. We also performed a metagenomic analysis of PT-containing microbes in fecal DNA using a novel NGS technology – PT-seq. With results revealing that 5-10% of gut microbes possess PTs, our studies lay the groundwork for future investigations of the role of PTs in IBD and other diseases.

## Results

### Optimizing the analytical and informatic platform for PT analysis in human fecal DNA

While PT modification gene systems are widespread among bacteria and archaea [13, 15, 16, 27] and there is evidence for PTs in human gut microbes [24], there has not been a systematic study of the true breadth, depth, and behavior of PT epigenetics in the gut microbiome. We initiated this exploration by developing a workflow (**Fig. 1A**) for identifying and quantifying PT dinucleotide spectra and PT-containing microbes in fecal DNA. Here we optimized existing technologies to improve yields of fecal DNA extraction (**Fig. 1B**) [28], to increase sensitivity and specificity of chromatography-coupled mass spectrometry (LC-MS) for quantifying PT dinucleotides [27], and to increase read depth for next generation sequencing (NGS)-based metagenomic analysis of the hundreds of genomes present in fecal DNA (**Fig. 1C**) [29]. The relatively low efficiency of fecal DNA extraction with the commercial Qiagen QIAmp Fast DNA Stool Mini Kit (**Supplementary Fig. S1A**) and the modest increase with the International Human Microbiome Standards (IHMS) Protocol Q [30, 31] (4-8 μg DNA/200 mg fecal material; **Supplementary Fig. S1A**), which extracted DNA from only 5% of a diluted sample, raised concerns about wasting samples and resources, while the harsh conditions used in Protocol Q raised concerns about PT stability during DNA isolation. To improve DNA yields, we revised the protocol by (1) avoiding the initial PBS dilution and homogenization steps and directly mixing fecal material with QIAamp InhibitEx buffer and (2) by increasing the concentration of fecal material in the dilution buffer. **Supplementary Figure S1B** shows that these changes increased the yield of DNA per mg of feces by more than 10-fold with a linear dose-response. The necessity of the bead-beating step is demonstrated in the 5-fold decrease in DNA yield in the absence of beads (**Supplementary Fig. S1C**). We further modified Protocol Q to increase the purity of the extracted DNA by adding an RNase A treatment to remove unwanted nucleic acid contamination and a second wash with buffer AW2 during the final cleanup steps before spin column elution. Overall, the optimized Protocol Q significantly outperformed the QIAmp Fast DNA Stool Kit in terms of DNA yield (**Supplementary Fig. S1D**), A_260_/A_230_ purity (**Supplementary Fig. S1E**), and A_260_/A_280_ purity (**Supplementary Fig. S1F**). Evidence for this improved purity is shown in **Supplementary Figure S1H,** with the optimized Protocol Q resulting in lower levels of UV-detectable contamination in fecal DNA digests resolved by HPLC.

**Fig. 1.**
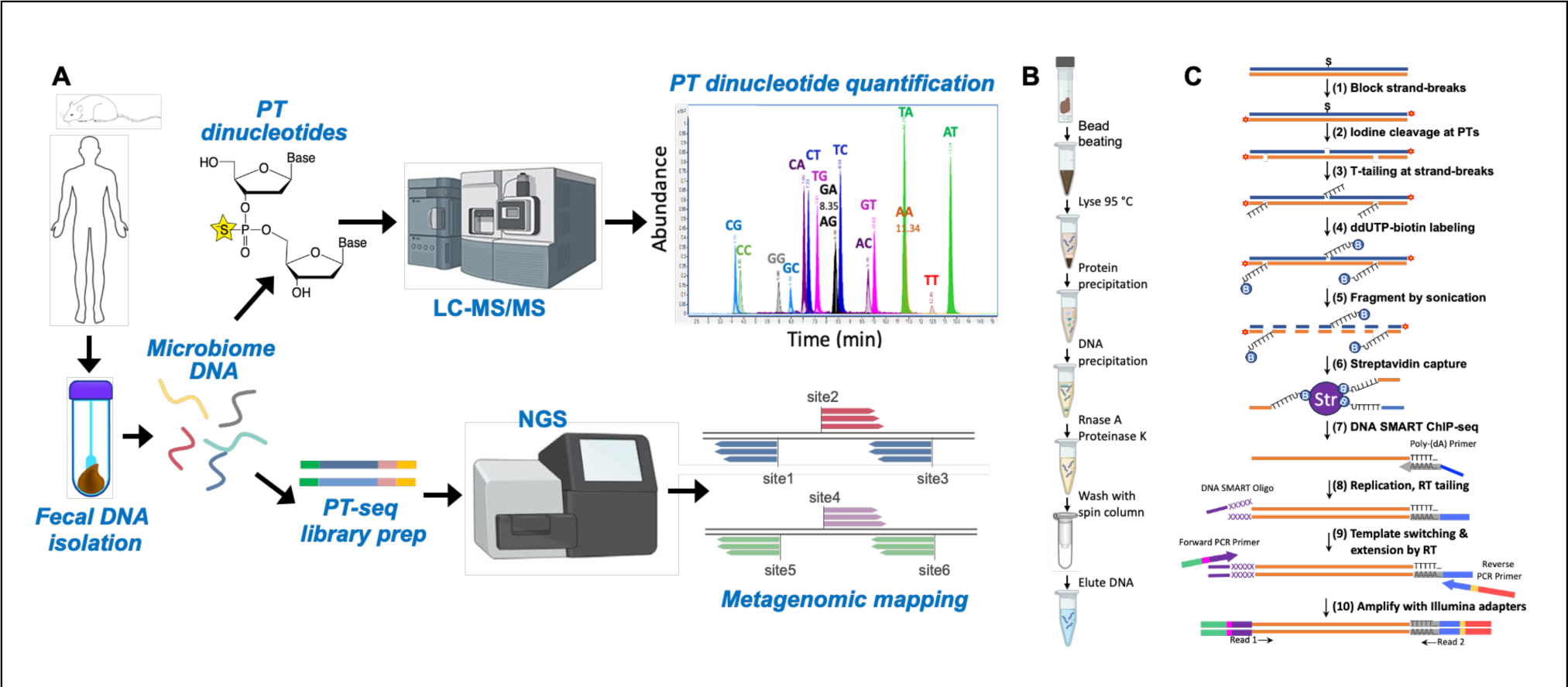
Workflows and optimized technologies for identifying and quantifying PT dinucleotides and mapping PT sites in gut microbes. (**A**) Purified fecal DNA was analyzed for PT dinucleotide content by LC-MS and subjected to PT-seq for metagenomic analysis of bacterial identity and PT consensus sequence. (**B**) Workflow for optimized fecal DNA isolation and purification, which improved yield by 10-fold. (**C**) Workflow for optimized PT-seq. The streptavidin capture step significantly reduced the level of noise and increased the read pileup sensitivity.

### A uniform spectrum of PT dinucleotides in the mouse gut microbiome

As a foundation for human studies, we first assessed the PT landscape in the mouse gut microbiome. With DNA extracted from fecal pellets obtained from individual mice, we quantified PT modifications as PT-bridged dinucleotides by exploiting the resistance of the backbone modification to nuclease P1 [12, 27], which results in a digest containing any of 16 possible PT dinucleotide combinations, all of which have the *Rp* stereochemical configuration of the phosphate [12]. The digestion mixture was analyzed by LC-MS with PT dinucleotides identified relative to chemical standards [12, 27]. The results of such an analysis are shown in **Figure 2A** (**Supplementary Table S1**). It is immediately apparent that the mice all shared a common spectrum of 11 of 16 possible dinucleotides in the same proportions, which might be expected for cage mates of a coprophagic mammal. The only significant difference (p < 0.02) between male and female mice involved dA_PT_dG (abbreviated A*G). Also telling was the quantity of each PT dinucleotide, which ranged from <1 to >90 per 10^6^ total nucleotides. This relatively low abundance compares to PT frequencies of ∼1 per 10^4^ in individual microbial genomes [27] and is most likely due to dilution of PT-containing bacteria by the other non-PT species present in the gut microbiome [3, 4]. However, this level of PTs suggests that an average of ∼5-10% of gut microbes possess PTs, which is consistent with the frequency of genes encoding PT synthesis proteins in >13,000 individual human microbiome isolates [17].

**Fig. 2.**
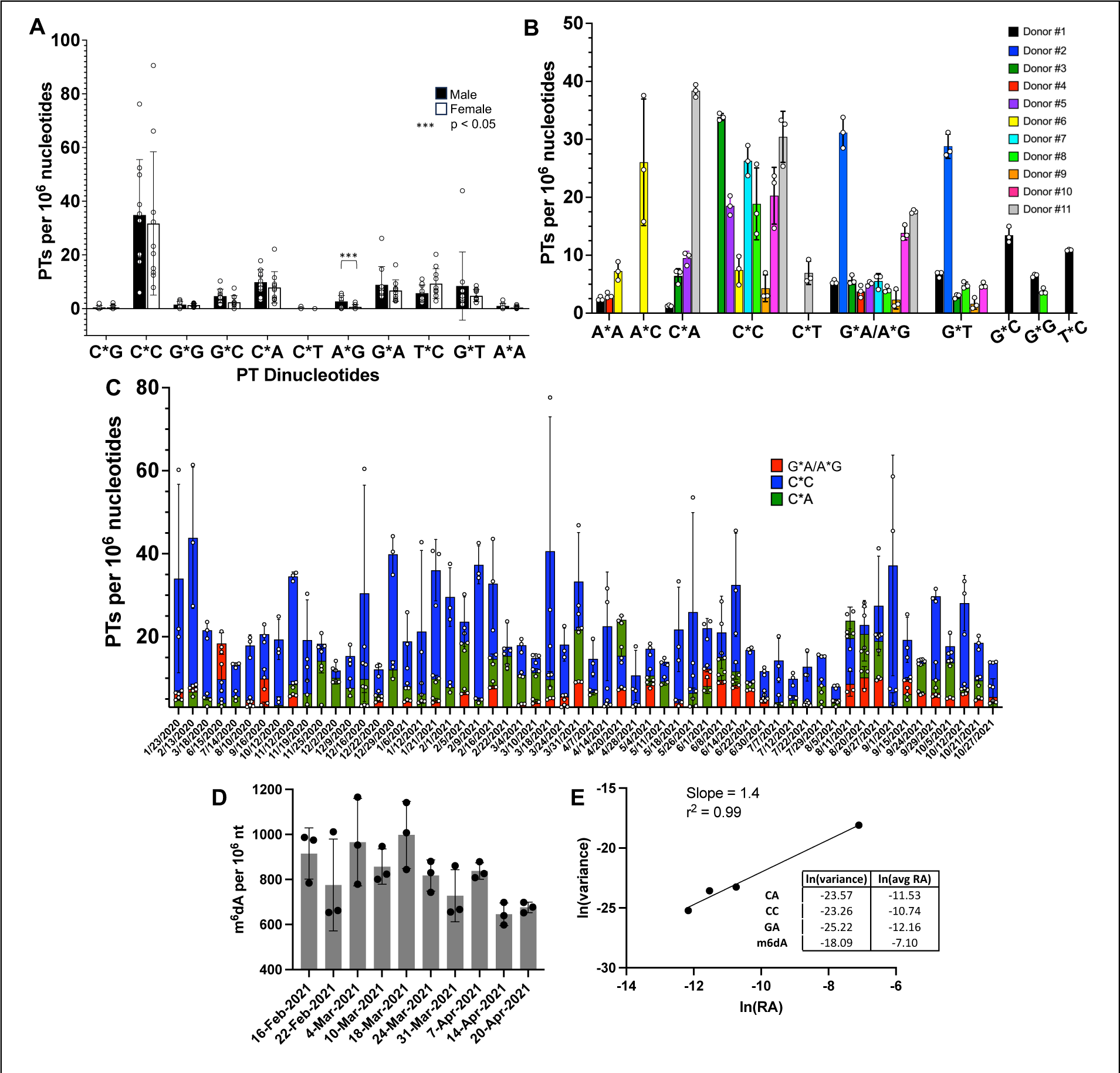
LC-MS analysis of mouse and human fecal DNA for PT dinucleotides by reveals the presence of PT-containing microbes. (**A**) Fecal DNA from individual C57BL/6 mice reveals a relatively uniform spectrum of PT dinucleotides with insignificant differences between males and females. Data represent mean ±SD for 20 mice. (**B**) Fecal PT dinucleotide spectra differ among 11 human donors. Data represent mean ±SD for N=11. (**C**) Analysis of PT dinucleotides in fecal DNA from donor #5 over 22 months reveals constant proportions of G*A, C*C, and C*A dinucleotides. Data represent mean ±SD for three separate DNA isolations from a single fecal sample. The bars representing PT levels for each of the 3 detected PT dinucleotides are superimposed, not stacked. (**D**) Overlay of m^6^dA levels on the time course of total PT dinucleotides from panel **C**. m^6^dA data represent mean ±SD for three separate DNA isolations from a single fecal sample. (**E**) Taylor’s Power Law (V = amb) analysis of temporal dynamics of PT dinucleotides and m6dA in Donor #5, where the mean abundance of a species (m) in a mixed population fluctuates over time with variance (V) linearly related to m to the power of b. Here a plot of ln(V) = ln(a) + b*(ln(m)) yields b = 1.4. (**F**) Correlations among the levels of the three PT dinucleotides from Donor #5.

### Diverse PT dinucleotide spectra in the human gut microbiome

We next compared the mouse data to humans. Analysis of fecal DNA from 11 healthy male and female donors revealed 10 of 16 possible PT dinucleotides (**Fig. 2B; Supplementary Fig. S2; Supplementary Table S2A**). Here we could not consistently chromatographically resolve A*G and G*A, so we combined the signals for both (denoted G*A/A*G). The results show significant diversity among the donors, with 2 to 7 different PT dinucleotides in each donor (**Fig. 2B**) and three PT contexts (G*A/A*G, G*T, and especially C*C) broadly shared among the donors. Among the dinucleotides we observed, the PT contexts G*A and G*T can derive from a known double-stranded sequence motif G*AAC/G*TTC modified by the *dnd* gene cluster in microbes such as *E. coli* B7A and *Salmonella enterica* serovar Cerro 8 [12, 27], while C*C was observed in the C*CA motif associated with the *ssp* gene cluster in *Vibrio cyclitrophicus* FF75 [19]. The 10 PT dinucleotide motifs detected here stand in contrast with the preliminary studies of Sun *et al*. [24] in which only 7 quantifiable PT dinucleotides were detected among 14 individuals; our analysis did not detect T*A but found 4 additional PTs (A*A, C*A, C*T, and G*C). These differences attest to the variation of microbiome PT epigenetics among humans. It is important to point out that the mass spectrometer signals for several PT dinucleotides were confounded by co-eluting non-PT species, which necessitated confirmation of the LC-MS behavior of all PT contexts for all 11 donors with exact mass analysis using a high-resolution mass spectrometer (**Supplementary Fig. S2**). This eliminated any uncertainty related to lower resolution triple-quadrupole analysis, chromatographic peak shifts, or low signal-to-noise ratios.

### Temporal dynamics of PT dinucleotides in the gut microbiome

Four donors contributed fecal samples regularly over periods of weeks to months to allow analysis of the temporal variation in levels of the PT dinucleotides in the gut microbiome. As shown in **Figure 2C** and **Supplementary Figures S4-S6 (Supplementary Tables S2B-S2F**), while the specific dinucleotide spectra remain roughly constant over time, the total level of all PT dinucleotides in each donor varies significantly on a weekly basis. For example, in donor #5, the presence of the C*A, C*C, G*A/A*G dinucleotides remains constant over the 21-month collection period while the level of total PTs varied by as much as 4-fold (**Fig. 2C**).

To correlate the PT fluctuations with other microbiome epigenetic marks, we measured levels of N^6^-methyl-2’-deoxyadenosine (m^6^dA), a widespread and abundant prokaryotic DNA modification [32, 33], in a subset of samples (10-weeks) from donor #5 and normalized the signals to levels of the canonical nucleosides, as with the PT dinucleotides (**Fig. 2D**; **Supplementary Table S3**). The presence of m^6^dA has been confirmed in lower eukaryotes [34], but its presence in mammalian cells is controversial [35] and, if real, would be orders-of-magnitude lower than the levels of 6-10 per 10^4^ nt observed here in fecal DNA (**Fig. 2D**) [35].

The magnitude of the temporal variation in PT levels in the four donors and the m^6^dA levels in Donor #5 raised the question of parallel behavior in the levels of PT-containing bacteria. The variation cannot be explained by time-dependent changes in PT levels in individual microbes. PT levels do not change significantly as a function of growth conditions [23, 36] likely due to their role in restriction-modification, with reduced PT levels in the face of unchanged restriction activity leading to high levels of DNA strand breaks and reduced fitness [37]. Here we tested the idea that fluctuations in the levels of microbiome DNA modifications followed Taylor’s power law, in which the mean abundance of a species in a mixed population (m) will fluctuate over time such that the variance (V) follows the equation V = am^b^. In human microbiomes, b is consistently >1, ranging roughly between 1.5 and 2 [38]. For the PT dinucleotides and m^6^dA levels in Donor #5, a plot of ln(V) = ln(a) + b*(ln(m)) yields b = 1.4 (**Figure 2E; Supplementary Tables S2B, S3, S4**), which is consistent with Ma’s definition of a Type III power law extension for mixed-species population spatial aggregation [38]. Further insights into possible shared niches of the various PT-containing microbes can be seen in the time course plots of individual PT dinucleotides for three donors in **Supplementary Figures S4-6**, in which changes in the levels of G*A/A*G and C*C are generally coordinated while those for G*T and G*G are not coordinated with each other or with G*A/A*G and C*C. The latter may thus be in similar growth environments or have similar dependencies. Similarly, the fluctuations in m^6^dA are apparently independent of the PTs. These behaviors beg the question of the identifies of PT-containing bacteria in the gut microbiome.

### Metagenomic analysis identifies PT-containing microbes and consensus sequences

To determine the identity of the microbes bearing PTs, we optimized our previously published methods for mapping PT sites using iodine to site-specifically cleave PTs and exploiting the strand breaks to locate PTs in genomes [19, 29]. While the methods proved useful for mapping PTs in pure populations of a single organism, such as the *Escherichia coli* B7A genome with its G*AAC/G*TTC consensus motif [19, 29], they were either too insensitive to analyze mixtures of hundreds of genomes as in fecal DNA or limited to bistranded PTs such as the G*AAC/G*TTC motif, thus ignoring single-strand PTs such as C*CA in *Vibrio cyclitrophicus* FF75 [19, 29]. Here we used an optimized PT mapping method that exploits the T-tailing approach [26, 29] to label iodine-nicked PT sites for subsequent extension, PCR amplification, and NGS sequencing, with attention to maximizing specificity and sensitivity for the complex mixture of genomes in the gut microbiome. With details provided in a separate publication [17], the general concept of “PT-seq” is outlined in **Figure 1C** and exploits iodine cleavage of PTs followed by 3’-end poly-(dT) tailing, ddUMP-biotin capping, and subsequent biotin-capture to enrich target DNA, and NGS sequencing library preparation by reverse transcriptase template switching. As a reference gut microbiome genome dataset for metagenomics analyses, we used both sequenced microbe isolates from the human gut in the Broad Institute-OpenBiome Microbiome Library [39, 40] and Metagenome-Assembled Genomes (MAGs) [41] to build a comprehensive custom collection of human gut microbiome (HGM) genomes (13,663 total) that reflected a global, healthy human gut microbiome population (see Methods). The data processing workflow for PT-seq and for parallel shotgun metagenomics of the gut microbiome is shown in **Supplementary Figure S7A**.

This approach was now applied to define the landscape of PT-containing microbes in fecal DNA from Donor #5. We first prepared a reference gut microbiome metagenome for this donor (**Supplementary Fig. S7B**) from an aliquot of isolated fecal DNA processed without 3’-end terminal blocking or iodine treatment. Taxonomical classification of these shotgun sequencing reads against the custom HGM collection, as well as additional reference databases (see Methods) to identify microorganisms not normally associated with the human gut microbiome (e.g., food), resulted in classification of 93% of the reads (**Supplementary Fig. S7B, Supplementary Table S5A**), leaving 7% of the reads as unclassified. Consistent with previously observed human gut microbiome compositions [5, 6], we found that Firmicutes was the most abundant phylum, followed by Bacteroidota, Actinobacteria, unclassified genomes, unclassified bacteria, Verrucomicrobia, and Proteobacteria (**Fig. 3A**, “Metagenome”). The number of reads assigned to each genome was used to evaluate relative abundance of the genome.

**Fig. 3.**
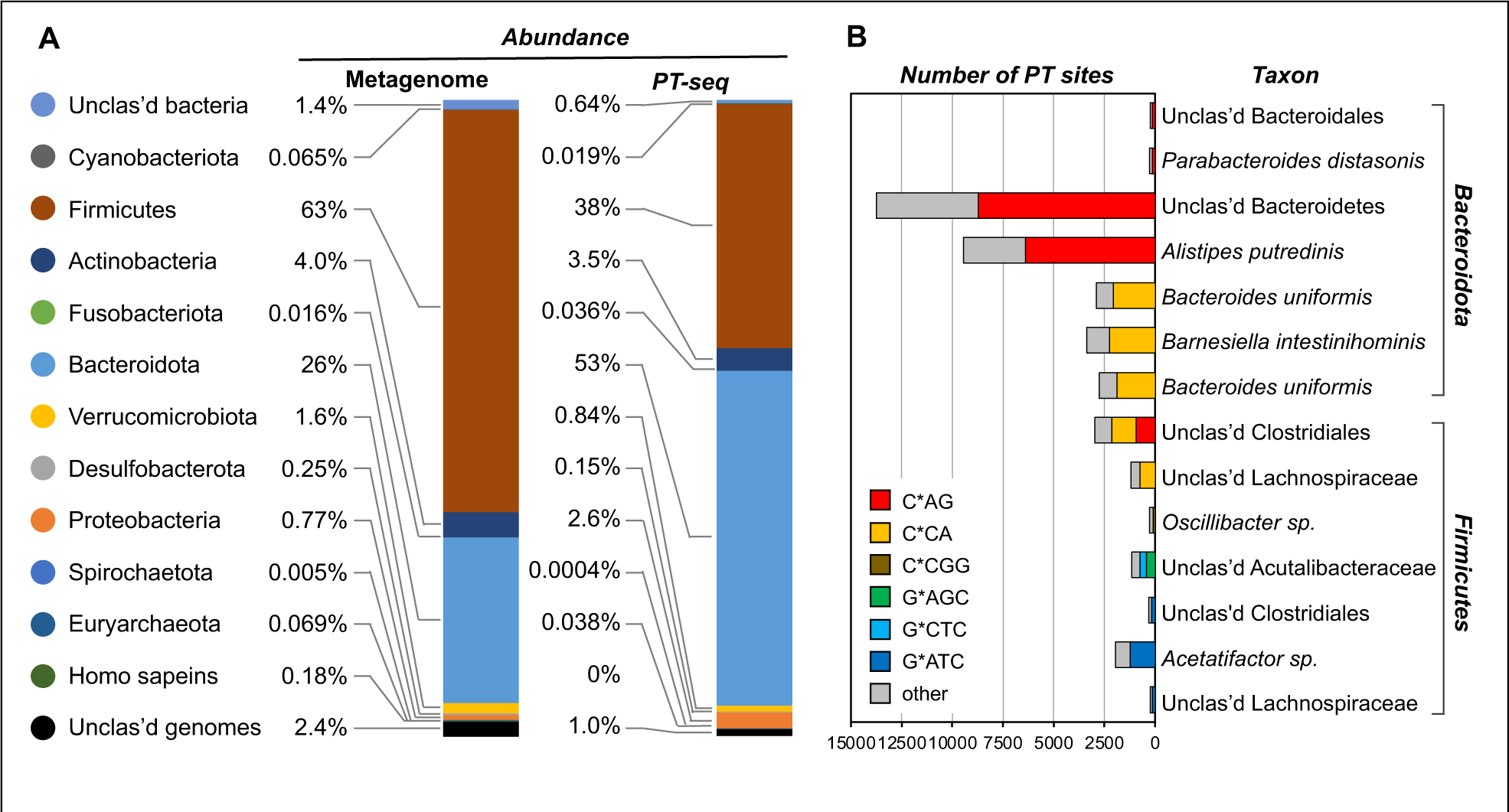
The taxonomic composition of microbiome and quantification of PT sites. (**A**) The abundance of microbiome was estimated by metagenomic sequencing using Kraken2 and Bracken. The phylogenetic composition of microbiome taxa collapsed at the phyla level in fecal sample (Donor #5). (**B**) The number of different PT modification motif sites identified by ICDS aligned to genomes of human gut microbiome isolates. A total of 26,817 sites were detected, with PTs denoted by a ‘*’.

We then applied PT-seq to another portion of isolated fecal DNA from Donor #5 and sequenced the library on an Illumina platform. The resulting reads (“PT-seq” in **Fig. 3A** and **Supplementary Fig. S7B**) were similarly cross-referenced against the custom HGM collection, resulting in assignment of 76% of PT-seq reads (**Supplementary Fig. S7B, Supplementary Table S5B**). We do not know the basis for the 17% difference in assigned reads for the metagenomic reference and PT-seq datasets, but it could be due to a greater proportion of small, unassignable DNA fragments resulting from the additional processing steps of PT-seq or greater resolution of contaminants due to biotin-capture enrichment. Bacteroidota emerged as the dominant phylum of PT-containing bacteria followed by Firmicutes, Actinobacteria, and Proteobacteria (**Fig. 3A**). These results indicated that the abundance of PT-containing bacteria in the gut microbiome did not simply mirror the taxonomic spectrum of gut bacteria in Donor #5.

### Correlating PT dinucleotides with metagenomic PT consensus sequences

To assign the PT-dinucleotides detected by LC-MS to specific gut microbes, we took advantage of the fact that PT-seq strand breaks occurred site-specifically at PTs and analyzed the sequences at the 5’-ends of the PT-seq reads to identify the consensus sequences of the PT-modifying enzymes in the PT-containing gut bacteria. Here we mapped PT reads to the top 100 genomes found in the HGM custom reference database using Mapper aligner software. The mapping results were further subjected to the Gaussian Mixture Models (GMM) to separate genomes by coverage, median sequencing depth, and dispersion of depth, which resulted in resolution of the 100 genomes into 3 clusters (**Supplementary Fig. S7C**). The first cluster contained 28 genomes with coverage >15% and with a larger median of sequencing depth, so they were likely to represent actual PT capture. Importantly, not all PT-seq sequences were necessarily relevant, since some alignments at this stage likely originated from contaminating DNA fragments from microorganisms that lack PT. In this regard, genomes in the other two clusters with <15% coverage, a smaller median of sequencing depth, and a larger dispersion were considered less likely to contain PT, since these could also arise from alignments among identical regions shared across many microorganisms.

Using the first cluster of 28 genomes, the consensus motif for PTs in each genome was determined by defining a span of 13-nt centered at each PT site in each read and subjecting these to MEME analysis [42] (**Supplementary Table S5C**). MEME detected five conserved motifs in 14 PT-containing genomes, including C*AG, C*CA, C*CGG, G*ATC, G*AGC/G*CTC, (**Fig. 3B, Supplementary Table S5D**). Among them, C*AG and C*CGG have not been reported in an organism previously. We recently confirmed the G*AGC/G*CTC motif in a human gut microbiome isolate *Lachnospiraceae sp.* (GMbC ID 2807EA_1118_063_H5) using PT-seq [17].

To validate these PT-seq-identified PT consensus sequences, we quantified read pileups along each of the 28 genomes with >15% coverage. Here we first established a minimal read depth for calling a read pileup as a PT site by adjusting the read depth from 1 to 25 and quantifying the total number of read pileups in the genome (**Supplementary Fig. S8**). As shown in **Supplementary Figure S8**, as the read pileup depth increased, the specificity for calling CA, CC, or GA sites, which were noted as sites of PT modification by LC-MS (**Fig. 2C**), among total pileup sites increased. Reaching a cutoff of 15 converged on 26,507 C*AG, C*CA, C*CGG, G*ATC and G*AGC sites that correlated with the three PT dinucleotides (C*A, C*C, and G*A) (**Fig. 2C**). This is roughly equivalent to 51 PTs per 10^6^ nucleotides, which is close to our LC-MS average of 43 ± 4 PTs per 10^6^ nt for the same fecal DNA sample analyzed by PT-seq (**Fig. 2C, Supplementary Data S2B)**. Therefore, we chose 15 as the optimal read pileup depth that balances sensitivity and specificity for calling sites of PT modification. Using this cutoff, we observed significant read pileups at 70% of putative PT modification sites in 2 out of 14 PT-containing genomes (**Supplementary Table S5D**). For example, the percentage of pileups at CAG sites was as high as 72.5% in an unclassified Clostridiales genome (GMbC ID 4452YQ_0918_057_F2), while the percentage of CCGG sites was as low as 38% in *Oscillibacter sp.* (genome ID GUT_GENOME141047) (**Supplementary Fig. S8**, **Supplementary Table S5D**), with these differences consistent with the frequencies of 3– and 4-nucleotide sequences in a genome, respectively. The observation of less than 100% calling of PTs at a putative modification sequence is consistent with the fact that PT modifications occur at only ∼10-15% of all possible consensus sequences in a bacterial genome [19]. Interestingly, despite not detecting G*C dinucleotides in our LC-MS analysis of Donor #5 fecal DNA, metagenomic analysis showed that GC sometimes increased in parallel with GA, especially in unclassified Acutalibacteraceae (genome ID GUT_GENOME238522), which would be consistent with a double-stranded G*AGC/G*CTC modification motif as observed in our recent publication [17]. In the genome of unclassified Clostridiales (GMbC ID 4452YQ_0918_057_F2), the specificity of CA and CC sites, which cannot be part of a short palindrome, increased simultaneously (**Supplementary Fig. S8, Fig. 3**), suggesting two distinct PT modification motifs in the same or similar microorganism(s). We observed 16,317 C*AG, 8,113 C*CA, 107 C*CGG, 1,543 G*ATC, 427 G*AGC and 310 G*CTC sites (**Fig. 3**, **Supplementary Table S5E**), which is equivalent to the following relative abundance: 33.5 C*A, 11.2 C*C, 6.8 G*A and 0.8 G*C. The lack of LC-MS detection of G*C is likely due to the five-fold lower sensitivity for detecting it compared to G*A (**Supplementary Figure S3**) and the relatively low level of G*A (**Fig. 2C**). This discrepancy points to the utility of metagenomics for discovering epigenetic modification sites and the complementarity of metagenomic and mass spectrometric analyses.

## Discussion

The gut microbiome provides an opportunity to characterize polymicrobial interactions in a community of hundreds of prokaryotic species and perhaps thousands of different bacteriophages [2–10]. While the importance of prokaryotic epigenetics is clear in microbial communities, for both defense and regulating gene expression, little is known about the role of microbial epigenetics in the dynamics of the human gut microbiome. Here we focused on chemically-reactive DNA phosphorothioate (PT) modifications as widespread prokaryotic and archaebacterial epigenetic marks [12, 19, 23, 27] with the potential for unique responses to inflammation [23].

Preliminary studies reported the presence of PTs in human fecal DNA [24] but lacked the technology to rigorously quantify the full spectrum of PTs and to identify PT-containing microbes. We thus set out to optimize, validate, and apply three technologies for defining the landscape and behavior of microbiome PT epigenetics (**Fig. 1**): fecal DNA extraction, LC-MS analysis of PT dinucleotides, and genome-wide site-specific mapping of PTs. For fecal DNA extraction, the 10-fold increase in DNA yield that we achieved not only accommodates the larger DNA needs of mass spectrometric analyses – an order-of-magnitude higher than NGS for metagenomics – but more importantly reduces biases introduced by highly diverse microbial cell wall mechanical properties and complex fecal matrices. Fecal DNA purified with this new method should more faithfully reflect the gut microbiome in health and disease. This increased fecal DNA fidelity combined with the optimized LC-MS method for PT dinucleotide quantification, which now minimizes matrix interferences, further increases the accurate and sensitive of detection of low-abundance PTs. There is still room for improvement in the chromatographic resolution and sensitivity for detecting the 16 PT dinucleotides, with nanoflow chromatography and ever-evolving mass spectrometry technology, such as ion mobility spectrometry [43]. The need for increased LC-MS sensitivity is illustrated by our observation of a PT-seq detection of a G*C-containing sequence in unclassified Clostridiales (**Fig. 3**) in Donor #5 despite the lack of LC-MS detection of G*C in the same fecal DNA (**Fig. 2**). As described in a companion publication [17], we revised and optimized previously published NGS PT mapping methods [19, 29] to develop PT-seq (**Fig. 1**), which allowed sensitive metagenomic analysis of PT-containing microbes and identification of PT consensus sequences.

Using these technologies, we found 10 PT dinucleotides shared by mice and humans (**Fig. 2**), while C*G was detected only in mice and A*C only in humans, thus accounting for 12 of 16 possible PT dinucleotides. The inability to detect C*G and A*C in their respective hosts or A*T, T*A, T*T, and T*G in either host may simply be due to low abundance of microbes bearing consensus sequences bearing these dinucleotide motifs. Indeed, if the detection of T*A by Sun *et al.* [24] is accurate, then only A*T, T*T, and T*G remain to be identified. The quantitative consistency of the PT dinucleotide spectra among individual male and female mice in our study was not unexpected given observations of highly consistent gut microbe populations in inbred mice due to genetics, shared environments, and coprophagy [44]. However, comparisons among different mouse cohorts in the MIT facility and between mice housed at other institutions will almost certainly reveal differences in the PT spectra in gut microbes given well established heterogeneity [44].

In the case of humans, the PT dinucleotide spectra were entirely unique for each of the 11 individual donors, with 2 to 7 of 10 different PT dinucleotides detected (**Fig. 2**). As PT-seq revealed, these dinucleotides are simply fragments of the 3-4 (or longer) consensus sequences for the Dnd, Ssp, and Brex epigenetic families, showing an enrichment in Bacteroidota, Firmicutes, Actinobacteria, and Proteobacteria (**Fig. 3**). These are also the major families of bacteria in the human microbiome, with our results shifting the balance from Firmicutes to Bacteroidota in the three seemingly independent groups of microbes with C*C, C*A, and G*A/A*G in Donor #5 (**Fig. 3**). Clearly we are analyzing the stool microbiome, which most resembles the luminal microbiome and likely does not sample the more controversial mucosal microbiome, especially the crypts [45]. Proteobacteria (Pseudomonodata) genuses Acinetobacter, Delftia, and Stenotrophomonas have been termed the “crypt-specific core microbiome” [46], but Bacteroidetes, Firmicutes, and nonfermentative Proteobacteria are also found in human colonic crypts [47]. It will be interesting to probe other gut microbiome compartments to assess the populations of PT-containing microbes.

We were initially perplexed by the temporal dynamics of PT dinucleotides and m^6^dA. One possible explanation is that PT levels vary significantly within individual microbes due to nutrient (e.g., sulfur) availability or stress. This possibility is ruled out, though, by the fact that bacteria maintain constant levels of PT throughout their genomes under a variety of growth conditions [23, 36] and the fact that reduced PT levels caused by deletion of synthesis genes leads to lethal DNA strand breaks by the remaining restriction system [37]. However, a quantitative test of Taylor’s Power Law revealed that the fluctuations simply reflected the normal dynamics of gut microbes, with less abundant microbes showing the smallest variance (**Fig. 2E**) [38]. The addition of the much more abundant m^6^dA increased the power of this analysis. The Taylor’s constant b = 1.4, which falls near the generic expectation of between 1.5 and 2, is entirely consistent with Ma’s definition of a Type III power law extension for temporal sampling of the microbiome [38]. It should be noted that none of the donors adhered to a special diet or strict lifestyle, suffered an illness, or used any antibiotics during the fecal collection period, so the PT compositions and time-dependent behaviors are likely due to the expected vague contributions of genetics [7] and environmental factors [8–10]. That the fluctuations in the PT levels obey expectation for gut microbes is consistent, once again, with the fact that the PT dinucleotides merely represent subsets of the microbiome population.

How do the levels of PT dinucleotides detected here compare to expectations based on the distribution of PT synthesis and restriction genes? The PT-seq results confirm subsets of the gut microbiome possess PT consensus sequences, but do not inform about frequency of those microbes. Jian *et al.* quantified PT-synthesis *dndA-E* genes in bacteria and archaea in the NCBI genome database and found *dnd* genes in 1.8% of bacteria and 0.7% of archaea, which is close to the 2.7% occurrence of the essential core set of *dndCD* in bacteria in our analysis of 13,000 gut microbiome isolates [17]. However, the occurrence level increases when other families of PT-synthesizing genes are considered, such as *ssp* and BREX, which accounted for an additional 5% of the gut microbe isolates [17]. This total of 7.7% of 13,000 gut microbiome isolates possessing PT-synthesis genes is consistent with the estimate of 5-10% of stool microbes possessing PTs based on (1) the quantity of PT dinucleotides at 1-90 per 10^6^ total nucleotides, (2) an average of ∼1 PT per 10^4^ nucleotides in individual microbial genomes [27], and (3) dilution of PT-containing bacteria by the other non-PT species present in the gut microbiome [3, 4].

## Conclusions

Here we optimized and applied fecal DNA extraction, LC-MS, and NGS PT-seq technologies to reveal a diverse landscape of PT epigenetics in 5-10% of human gut microbes. Within the context of 11 healthy human fecal donors, LC-MS analysis of limit digests of fecal DNA revealed signature combinations and proportions of the 16 possible PT dinucleotides that reflect bacterial restriction-modification consensus sequences. The PT dinucleotides displayed temporal dynamics consistent with Taylor’s Power Law and the contribution of their host bacteria to the gut population. Application of PT-seq for site-specific metagenomic analysis of PT-containing bacteria in one fecal donor revealed the larger consensus sequences for the PT dinucleotides in Bacteroidota, Firmicutes, Actinobacteria, and Proteobacteria, and further revealed an additional PT consensus sequence not detected by LC-MS. Our results provide a benchmark for understanding the behavior of an abundant and chemically-reactive epigenetic mark in the human gut microbiome, with implications for inflammatory conditions of the gut.

## Methods

### Materials

Nuclease P1 (N7000) was purchased from US Biological, Inc and reconstituted according to the manufacturer’s protocol. Calf intestinal alkaline phosphatase (P5521) was from Sigma and reconstituted according to the manufacturer’s protocol. Spin filters (10 kDa) (82031-350) were from VWR. The QIAmp Fast DNA Stool Mini Kit (51604) was purchased from Qiagen. The PowerLyzer PowerSoil (12855) kit was purchased from MO BIO Laboratories, Inc., now part of Qiagen. RNase A (12091021) was purchased from Invitrogen. Ammonium acetate (7.5 M) (A2706) was purchased from Sigma.

### Mouse fecal collection, DNA isolation, and LC-MS analysis of PT dinucleotides

Healthy C57BL/6 mice from The Jackson Laboratory in Bar Harbor, ME were 31-32 days old at the time of collection and weighed from 16g – 23g. Fecal pellets (3-4) were collected from individual male (10) and female (10) mice under an approved protocol (MIT Committee on Animal Care Protocol 0912-093-15) by removing mice from cages, placing them on a clean bench surface, and collecting freshly excreted fecal pellets. Fresh pellets were placed in a 1.5 mL Eppendorf tube for each mouse and immediately frozen on dry ice, followed by long-term storage at –125 °C. DNA was isolated form the pellets using the PowerLyzer PowerSoil kit according to manufacturer instructions and all isolations were performed on the same day to reduce experimental bias. Isolated DNA was concentrated under vacuum (SpeedVac, Savant, ThermoFisher), its concentration determined by Nanodrop (ThermoFisher), and the concentration adjusted to 50 ng/μL. For LC-MS/MS analysis, 50 μL (∼2.5 μg) of DNA were digested with 0.5 U Nuclease P1 (50 °C, 60 min.), followed by 1U calf intestine phosphatase for 1 h at 37 °C. Samples were filtered with a VWR 10kD MWCO filter to remove proteins and concentrated under vacuum to ∼ 30 μL.

PT-linked dinucleotides in the DNA digest (∼1µg) were analyzed by LC-MS/MS on a system consisting of an Agilent 6430 ESI-QQQ coupled with an Agilent 1290 HPLC equipped with a diode-array detector (DAD) set to 260 nm absorbance. Chromatographic separation of PT dinucleotides was achieved using a Phenomenex Fusion-Rp column (2.5 µm particle size, 100 Å pore size, 100 mm length, 2 mm i.d.) eluted with a gradient running from 97% buffer A (5 mM NH_4_OAc, pH 5.3) to 9% buffer B (acetonitrile) over 13 min followed by a 1 min column wash with 95% B and re-equilibration with 97% buffer A for 3 min. The QQQ was operated with dynamic multiple reaction monitoring (DMRM) in positive ion mode with the following source parameters: N_2_ gas temperature 350 °C and 10 L/min flow rate, nebulizer pressure 40 psi and capillary voltage 3500 V. HPLC retention times and mass transitions for each of the 16 PT-linked dinucleotides were as detailed in previous publications [12, 23, 27], similar to **Table 1** below. Response factors for absolute PT quantification were calculated using external calibration standard curves based on synthetic dinucleotide standards prepared as described previously [12, 23, 27]. Standards were also run in parallel with samples to confirm retention times.

**Table 1.** Mass spectrometry parameters for 16 PT dinucleotides and canonical 2’-deoxyribonucleotides.

| **Compound** | **Exact Mass** | **Precursor Ion** | **Product Ion** | **Retention Time (min)** | **Collision Energy** | **Cell Accelerator Voltage** |
| --- | --- | --- | --- | --- | --- | --- |
| d(A*A) | 581.14388 | 581.1 | 136.1 | 12.4-12.5 | 40 | 3 |
| d(A*C) | 557.13264 | 557.1 | 136.1 | 10.9-11.0 | 32 | 3 |
|  |  | 557.1 | 112.1 | 10.9-11.0 | 24 | 3 |
| d(A*G) | 597.13879 | 597.1 | 152.1 | 9.61-9.70 | 48 | 5 |
|  |  | 597.1 | 136.1 | 9.70 | 36 | 1 |
| d(A*T) | 573.13231 | 572.1 | 136.1 | 14.8 | 32 | 1 |
|  |  | 572.1 | 127.1 | 14.8 | 48 | 5 |
| d(C*A) | 557.13264 | 557.1 | 136.1 | 8.27-8.29 | 32 | 1 |
|  |  | 557.1 | 112.1 | 8.27-8.29 | 28 | 3 |
| d(C*C) | 533.12141 | 533.1 | 112.1 | 5.31-5.47 | 16 | 5 |
| d(C*G) | 573.12756 | 573.1 | 152.1 | 5.11-5.31 | 32 | 1 |
|  |  | 573.1 | 112.1 | 5.11-5.31 | 32 | 5 |
| d(C*T) | 548.12108 | 548.1 | 127.1 | 8.79-8.81 | 36 | 5 |
|  |  | 548.1 | 112.1 | 8.79-8.81 | 28 | 5 |
| d(G*A) | 597.13879 | 597.1 | 152.1 | 9.61-9.70 | 28 | 3 |
|  |  | 597.1 | 136.1 | 9.61-9.70 | 32 | 1 |
| d(G*C) | 573.12756 | 573.1 | 152.1 | 7.64-7.65 | 32 | 5 |
|  |  | 573.1 | 112.1 | 7.64-7.65 | 40 | 3 |
| d(G*G) | 613.13371 | 613.1 | 152.1 | 7.14-7.20 | 36 | 3 |
| d(G*T) | 588.12722 | 588.1 | 152.1 | 11.5-11.6 | 32 | 5 |
|  |  | 588.1 | 127.1 | 11.5-11.6 | 40 | 3 |
| d(T*A) | 573.13231 | 572.1 | 136.1 | 12.92 | 28 | 3 |
|  |  | 572.1 | 127.1 | 12.9 | 44 | 3 |
| d(T*C) | 548.12108 | 548.1 | 127.1 | 10.1-10.2 | 44 | 1 |
|  |  | 548.1 | 112.1 | 10.1-10.2 | 24 | 5 |
| d(T*G) | 588.12722 | 588.1 | 152.1 | 9.18-9.20 | 32 | 5 |
|  |  | 588.1 | 127.1 | 9.18-9.20 | 48 | 1 |
| d(T*T) | 563.12074 | 563.1 | 127.1 | 14.4 | 40 | 5 |
| dA | 252.10912 | 252.1 | 136.1 | 5.23 | 12 | 1 |
| dC | 228.09788 | 228.1 | 112.1 | 1.45 | 16 | 1 |
| dG | 268.10403 | 268.1 | 152.1 | 2.71 | 4 | 3 |
| dT | 243.09755 | 243.1 | 127.1 | 3.22 | 12 | 1 |

### Collection of human feces

Collection of human fecal matter from healthy adult volunteers was performed under a protocol approved by the MIT Committee on Use of Human Subjects (Protocol 2306001007). Donors were provided with a kit consisting of a large weigh boat to assist collection, a plastic micro spatula for manipulation, a 50 ml centrifuge tube for storage, a pair of large nitrile gloves, a re-sealable zipper plastic bag for organization and temporary ice storage, and a small paper bag for concealment. Donors were instructed to drop off fecal samples for processing immediately, or place the sample at 4 °C for up to 3 h prior to delivery, or store the samples at –20°C. All samples were processed within 24 h of collection in any event.

### Optimization of Protocol Q for DNA isolation from feces

To enhance the yield of DNA from fecal extraction and ensure an unbiased population of microbial genomes, we optimized each step of the International Human Microbiome Standards (IHMS) IHMS_SOP 06 V3: Standard operating procedure for fecal samples DNA extraction, Protocol Q [28].

#### Amount of human fecal material

Varying amounts of fecal material were weighed into a 15 mL tube (200, 400, or 800 mg). To the tube was added 4 mL of sterile PBS (10 mM, pH 7.4). The mixture was homogenized using a Qiagen tissue homogenizer at full speed for 1 minute. 200 µL of this mixture was used for the DNA isolation and the isolation procedure proceeded as published in IHMS Protocol Q (IHMS_SOP 06_v2 from 4/12/2015).

#### Bead beating step

Varying amounts of fecal material were weighed into a 2 mL tube (100, 150, or 200 mg). Samples were prepared in duplicate, one set to be processed with beads and one set to be processed without beads. The DNA isolation followed IHMS Protocol Q as published in IHMS_SOP 06_v2 from 4/12/2015 with several modifications: After the addition of 1 mL of InhibitEx buffer to the 2 mL tube containing fecal material, the sample was homogenized on a FastPrep homogenizer for a total of 2 cycles, 45 seconds per cycle at a speed setting of 6.0.

#### PT dinucleotide stability following timed heating at 95°C

Here we added 20 µg (200 µL) of purified genomic DNA isolated from *E. coli B7A* to a 2 mL tube. Our DNA isolation protocol was carried out exactly as described above but with varying incubation times at 95 °C. Heating was carried out for either 5, 10, 15, or 20 min. After completion of the protocol, the DNA was digested to a mixture of mononucleosides and PT-containing dinucleotides using NP1 and CIAP as described above. PT dinucleotide levels were quantified by LC-MS as described above.

### Extraction of bacterial DNA from human fecal samples using optimized Protocol Q

Fecal material (180-220 mg) was combined with 0.1 mm (0.3 g) and 0.5 mm (0.3 g) Zirconia beads and 1 mL of InhibitEx buffer from the QIAmp Fast DNA Stool Mini Kit in a 2 mL tube. The mixture was processed on a FastPrep Cell Disruptor for 45 s using a speed setting of 6.0 (2 total cycles of bead beating). The mixture was then incubated at 95 °C for 15 min. The mixture was again homogenized on a FastPrep Cell Disruptor for 45 seconds using a speed setting of 6.0 (2 total cycles of bead beating). Solid material was pelleted via centrifugation at 4 °C for 5 min. at 16,100 g. The supernatant was transferred to a new 2 mL tube and kept on ice. InhibitEx buffer (300 µL) was added to the pellet and the mixture was again homogenized on a FastPrep Cell Disruptor for 45 seconds using a speed setting of 6.0 (2 total cycles of bead beating). Solid material was pelleted via centrifugation at 4 °C for 5 min. at 16,100 g. The supernatant was pooled with the previous supernatant. Ammonium acetate (260 µL of 7.5 M) was added to the pooled supernatants, and the solution was vortexed and incubated on ice for 5 min. Precipitate was pelleted by centrifugation at 4 °C for 10 min. at 16,100 g. The supernatant was removed and aliquoted into separate 2 mL tubes (650 µL each). One volume (650 µL) of isopropanol was added to each aliquot, the mixtures were vortexed, and incubated on ice for 30 min. DNA was pelleted by centrifugation at 4 °C for 15 min. at 16,100 g. The supernatant was discarded and pellets were gently washed with 70% ethanol (500 µL) by inverting 3 times before centrifugation at 16,100 g at 4 °C for 15 min. Excess supernatant was removed by aspiration, pellets were air dried for 10 min, and then solubilized in 100 µL Tris-HCl (10 mM, pH 8.0). The solutions were pooled, combined with RNase A (2 µL of 20 mg/mL) and incubated at 37 °C for 10 min.

Proteinase K (15 μL) was added to a 1.5 mL tube before the addition of the resuspended precipitated DNA (200 μL) and Buffer AL (200 μL) from the Qiagen Fast DNA Stool Kit. The mixture was vortexed (15 s) and incubated at 70 °C for 10 min. Ethanol (200 μL) was added to the mixture and the entire mixture (600 μL) was added to a QIAmp spin column, centrifuged at 16,100 g for 1.5 min, and the flow-through was discarded. The column was then washed with Buffer AW1 (600 μL), centrifuged at 16,100 g for 1.5 min, and the flow-through was discarded. The column was then washed twice with Buffer AW2 (600 μL) as above. The column was centrifuged at 16,100 g for 3 min to remove residual buffer. Pre-heated Tris-HCl (100 μL, 10 mM, pH 8.0) was applied to the column, incubated for 1 min, and centrifuged at 16,100 g for 1.5 min to elute the DNA. The yield and purity of DNA was assessed by Nanodrop. DNA was stored at –20 °C.

### Digestion of DNA for LC-MS/MS analysis of PT dinucleotides

DNA (20 μg, 78 μL) was incubated with Nuclease P1 (1.5 U, 3 μL) in 30 mM ammonium acetate pH 5.3 and 0.5 mM ZnCl_2_ (90 μL total reaction volume) for 2 h at 55 °C. The reaction mixture was diluted with Tris-HCl (∼100 mM final concentration, pH 8.0, 9 μL) and incubated with calf intestinal alkaline phosphatase (51 U, 3 μL) for 2 h at 37 °C. Enzymes were removed by passing the mixture through a VWR 10 kDa spin filter with centrifugation at 12,000 g for 12 min. The solution was lyophilized to dry and resuspended in H_2_O (50 μL).

### LC-MS/MS analysis of PT dinucleotides

Synthetic PT DNA dinucleotides or Nuclease P1 hydrolyzed DNA were analyzed by LC-MS/MS on an Agilent 1290 series HPLC system equipped with a Synergi Fusion RP column (2.5 μm particle size, 100 Å pore size, 100 mm length, 2 mm inner diameter) and a DAD. The HPLC was coupled to an Agilent 6490 triple quadrupole mass spectrometer. The column was eluted at 0.35 mL/min at 35 °C with a linear gradient of 3-9% solvent B (acetonitrile) in solvent A (5 mM ammonium acetate pH 5.3) over 15 min. The column was then washed with 95% solvent B for 1 min, and then re-equilibrated with 97% solvent A for 3 min. Canonical deoxyribonucleosides that eluted from the column were quantified by their 260 nm absorbance with the DAD. PT-containing dinucleotides were identified and quantified by tandem quadrupole mass spectrometry with electrospray ionization operated with the following parameters: N_2_ temperature, 200 °C; N_2_ flow rate, 14 L/min; nebulizer pressure, 20 psi; capillary voltage, 1800 V; and a fixed fragmentor voltage, 380 V. For product identification, the mass spectrometer was operated in positive ion polarity using dynamic multiple reaction monitoring (DMRM) mode with the conditions tabulated below.

### High resolution mass spectrometry

Synthetic PT dinucleotides (2 pmol per 10 μL injection) or Nuclease P1 hydrolyzed RNA (4 μg per 10 μL injection) were analyzed on a Dionex Ultimate 3000 UHPLC system equipped with a Synergi Fusion RP column (2.5 μm particle size, 100 Å pore size, 100 mm length, 2 mm inner diameter). The HPLC was coupled to a Thermo Fisher Q Exactive Hybrid Quadrupole-Orbitrap mass spectrometer. The column was eluted at 0.35 mL/min at 35 °C with a linear gradient of 3-9% acetonitrile in 97% solvent A (5 mM ammonium acetate pH 5.3) over 15 min. The column was washed with 95% acetonitrile in solvent A for 1 min, and initial conditions were regenerated by equilibrating the column with 97% solvent A for 3 min. High resolution mass spectra for the PT-containing dinucleotides were obtained by hybrid quadrupole-Orbitrap mass spectrometry with the following parameters: sheath gas flow rate, 50 L/min; aux gas flow rate, 15 L/min; sweep gas flow rate, 3 L/min; spray voltage, 4.20 kV; and capillary temperature, 275 °C. For product identification, the mass spectrometer was operated in positive ion polarity using targeted single ion monitoring mode with the conditions tabulated in Table 1.

### Raw mass spectrometry data processing and statistical analysis

Raw data were processed using either Agilent MassHunter Qualitative Analysis software for samples analyzed on the Agilent 6490 instrument or Thermo Xcalibur Qual Browser for samples run on the QExactive instument. Only signals with distinct peak shape, determined by visual inspection, were considered for data-analysis. Each PT dinucleotide signal was correlated to a response factor from runs using calibration standards. Absolute PT levels per 10^6^ nucleotides were calculated based on the estimated amount of injected canonical nucleosides using UV signals at 260 nm in correlation with UV calibration curves.

### Metagenomic sequencing

For shotgun metagenomics, 5 µg DNA was extracted from a Donor #5 fecal sample using optimized protocol Q and submitted to Novogene (UC Davis, California). After QC, DNA was fragmented and subjected to end repair and phosphorylation. Next, A-tailing and adaptor ligation was performed in a PCR-free method. Finally, 150 bp paired-end sequencing was performed with the Illumina Novaseq 6000.

### Defining reference genomes for metagenomic sequencing

The first step in mapping PT modifications was to define a reference human genome for comparisons with our human gut microbiome sequences, using both microbe isolates and Metagenome-Assembled Genomes (MAGs). We designed a comprehensive custom collection of human gut microbiome (HGM) genomes that reflect a global, healthy human gut microbiome population. This reference collection contains 3,632 genomes of gut bacterial isolates from the Broad Institute-OpenBiome Microbiome Library (BIO-ML) [39], 5387 genomes of gut bacterial isolates from the Global Microbiome Conservancy (GMbC) collection [40], and 4,644 genomes of gut microbal isolates and MAGs from the Unified Human Gastrointestinal Genome (UHGG) collection [41]. The 9019 genomes from GMbC and BIO-ML were filtered using drep with 99% average nucleotide identity[48]. We taxonomically classified sequencing reads against the custom reference database (50,67 genomes total) using Kraken2 [49] and Bracken [50].

### Library preparation for PT-seq

As detailed in an accompanying paper, optimized signals for PT-seq were achieved in part by blocking of pre-existing strand-break sites was achieved by 4 cycles of denaturation-dephosphorylation-blocking. Each cycle started by denaturing at 94 °C for 2 min and immediately cooling down on ice for 2 min. The initial dephosphorylation reaction was in a mixture (50 μL) containing 5 μL of terminal transferase buffer (NEB Tdt Reaction Buffer, Catalog # M0315S), 1 μL of shrimp alkaline phosphatase (rSAP, NEB Catalog # M0371S), and 10 μg of fragmented DNA, with incubation at 37 °C for 30 min to remove phosphate at 3′ end of the strand-breaks. The phosphatase was then inactivated by heating at 65 °C for 10 min. After cooling, 1 μL of Tdt Reaction Buffer, 6 μL of CoCl_2_ (0.25 mM), 2 μL of ddNTPs (2 mM each, TriLink) and 1 μL of terminal transferase (20 units, NEB Catalog #M0315S) was added to the reaction with incubation at 37 °C for 1 h to block any pre-existing strand-break sites. Blocking cycles were repeated 3 times, in which fresh reagents were added in each cycle (0.25 μL of Tdt Reaction Buffer, 0.25 μL of CoCl_2_, 2 μL of ddNTPs and 0.5 μL of Tdt). The blocked DNA was purified using a DNA cleanup kit (Zymo Catalog # 11-304C).

For iodine cleavage, 32 μL of the blocked DNA was incubated with 4 μL of 500 mM Tris-HCl pH 10.0 and 4 μL of iodine solution (50 mM, Fluka, Catalog # 318981-100) at room temperature for 5 min. Then, the reaction product was purified using two DyeEX columns (QIAGEN Catalog #63206) to remove salts and iodine. The purified DNA was denatured by incubating at 94 °C for 2 min and cooling on ice for 2 min and then incubated with 4 μl of H_2_O, 5 μL of NEB rCutsmart buffer and 1 μL of rSAP (50 μl reaction) to remove 3′-phosphates arising from iodine cleavage. After incubation at 37 °C for 30 min and 65 °C for another 10 min, the product was incubated with 1 μl of dTTP (1 mM, NEB Catalog # N443S), 1 μL of Tdt buffer, 6 μL of CoCl_2_, 1 μL of Tdt and 1 μL of H_2_O. After incubation at 37 °C for 45 min, the dTTP was removed by DyeEX columns.

DNA (60 μL) was then incubated with 7.8 uL of Tdt buffer, 7.8 uL of CoCl_2_, 1 μL of ddUTP-biotin (1 mM, Jena Bioscience NU-1619-BIOX-S), 1 μL of Tdt and 1 μL of H_2_O at 37 °C for 1 h to terminate the T-tails by ddUTP-biotin. After cleaned with DyeEX columns, the DNA was diluted in 500 μL of H_2_O and fragmented by probe sonication as described above. The DNA fragments was mixed with 10 μL of streptavidin-coated magnetic beads (NEB Catalog # S1421S) and 500 μL of binding buffer (5 mM Tris-HCl, pH 7.5, 1 M NaCl, 0.5 mM EDTA) and incubated on a shaker at ambient temperature for 1 h. The beads were pull down by a magnetic and washed 3-times with 100 μL of binding buffer. After discarding the supernatant, the beads were resuspended in 20 μL of H_2_O.

The purified product in ICDS method and captured beads in upgraded ICDS method were ready for Illumina library preparation with SMART ChIp-seq kit (Takara, Catalog # 634865) by following the manufacturer’s protocol. The final step of PCR was performed using the Illumina primers provided in ChIp-seq kit and 12 cycles were used for amplification. The PCR product with unique sequencing barcode was submitted to Illumina MiSeq and NovaSeq instrument for 150 bp paired-end sequencing.

### Data analysis

Adapters were removed using bbduk from BBtools (sourceforge.net/projects/bbmap/), (with parameters ktrim=r k=18 hdist=2 hdist2=1 rcomp=f mink=8 qtrim=r trimq=30 for R1, ktrim=r k=18 mink=8 hdist=1 rcomp=f qtrim=r trimq=30 for R2). T-tails were removed using bbduk with parameters ktrim=r k=15 hdist=1 rcomp=f mink=8 for R1 and ktrim=l k=15 hdist=1 rcomp=f mink=8 for R2. PhiX were removed using bbduk with parameters k=31 hdist=1.

We collected 3,632 genome sequences from the BIOML[39] and 5,387 genome sequences from GMbC [40]. We filtered genomes at an estimated species level (ANI≥99%) using dRep v2.5.4 [48] with the options: –sa 0.99. We collected 4,644 genome sequences from the UHGG collection [41]. For metagenomic sequencing data, the trimmed reads were classified against the custom library built with filtered human gut microbiome sequences from BIOML, GMbC and UHGG using Kraken2 [49] and Bracken [50]. The relative abundances of genomes were calculated using the number of reads assigned.

For PT-seq, we mapped the resulting reads to 100 genomes with the most reads assigned using Mapper v1.1-beta04 (github.com/mathjeff/Mapper). If a read maps to multiple positions with the optimal alignment quality, we output all alignment results and computed depth to each position as 1/n (n=number of optimal aligned positions). The coverage and depth were subjected to clustering using the Gaussian Mixture Models (GMM). Twenty-eight genomes in the cluster with more than 15% coverage were then subjected to consensus motif detection. Using custom scripts, the 13-nt sequences centered at 5’-ends of read pileups were retrieved from vcf files generated by Mapper. With incrementing cutoff of depth of read pileups from 1 to 25, the number of pileups at each NN nucleotide site were counted using homemade scripts and the motifs were analyzed using MEME [42] with parameters: –dna –objfun classic –nmotifs 5 –mod zoops –evt 0.05 –minw 3 –maxw 6 –markov_order 0 –nostatus –oc. The equivalent PTs were calculated by the number of read pileups derived from the genome size and normalized to the relative abundance.

## Declarations

### Ethics approval and consent to participate

Mouse studies were performed under a protocol approved by the MIT Committee on Animal Care (Protocol 0912-093-15). Human studies were performed under a protocol approved by the MIT Committee on Use of Human Subjects (Protocol 2306001007). As there is no identifying information associated with the data presented here, consent for participation and publication was not required.

## Consent for publication

## Availability of data and material

Raw PT-seq data has been uploaded to the NCBI SRA database with BioSample ID SAMN39420940. Raw LC-MS data have been deposited to the ProteomeXchange Consortium via the PRIDE [51] partner repository with the dataset identifier PXD051389. During the peer review process data will be accessible using the following details: Username: Password: gZWilfPq

## Acknowledgements

Financial support was provided by a NIEHS Training Grant in Environmental Toxicology T32-ES007020 (S.R.B.), Core Center Grant P30-ES002109 from the NIEHS (P.C.D.), from the Deutsche Forschungsgemeinschaft (S.K.), and from an NIH Transformative Award # R01-OD028099-01 (P.C.D., E.J.A.).

## Author Contributions

S.R.B., M.S.D., Y.Y., J.G.F., E.J.A., and P.C.D. planned the study. S.R.B., M.S.D., and Y.Y. directly performed most experiments. F.G. analyzed time course data from donor #5 for periodicity shown in Fig. S14. S.K. performed the mouse experiments and analyzed the data described in Fig. 3. S.R.B., M.S.D., Y.Y., and P.C.D. performed data analysis. S.R.B, M.S.D., Y.Y., J.G.F., E.J.A., and P.C.D. wrote the manuscript.

## Supplementary Information for

## Supplementary Figures

**Fig. S1.** Optimization of Protocol Q for fecal DNA extraction.

**Fig. S2.** Exact mass confirmation of PT dinucleotides using Orbitrap for donors #1-#11.

**Fig. S3**. LC-MS calibration curves for PT dinucleotides for donor #5.

**Fig. S4.** PT dinucleotide time course for donor #7 for **(A)** total PTs, **(B)** G*A, and **(C) C*C**.

**Fig. S5.** PT dinucleotide time course for donor #8 for (**A**) total PTs, (**B**) G*T, (**C**) G*G, (**D**) G*A, and (**E**) C*C.

**Fig. S6**. PT dinucleotide time course for donor #9 for (**A**) total PTs, (**B**) G*T, (**C**) G*A, and (**D**) C*C.

**Fig. S7**. Workflow for processing of metagenomic sequencing and PT-seq data.

**Fig. S8**. The number of read pileups (y-axis) with an increasing depth cutoff (x-axis) in 28 genomes with >15% coverage.

**Fig. S9**. The percentage of read pileups at each of 16 possible dinucleotide sites.

## Supplementary Tables – all as separate Excel spreadsheets

**Table S1.** PT dinucleotides in fecal DNA from mice. This source data is associated with **Figure 2A**.

**Table S2A**. PT dinucleotides in fecal DNA from 11 human donors. This source data is associated with **Figure 2B**.

**Table S2B**. Time course of PT dinucleotide levels in Donor #5. This source data is associated with **Figure 2C, 2D, 2E**.

**Table S2C**. Time course of PT dinucleotide levels in Donor #7. This source data is associated with **Figure S4**

**Table S2D**. Time course of PT dinucleotide levels in Donor #8. This source data is associated with **Figure S5**.

**Table S2E**. Time course of PT dinucleotide levels in Donor #9. This source data is associated with **Figure S6**.

**Table S2F**. Calibration curves for PT dinucleotides and canonical nucleosides. This source data is associated with **Figure S3**.

**Table S3**. Time course of m^6^dA in Donor #5. This source data is associated with **Figures 2C** and **2D**.

**Table S4.** Taylor’s power law analysis of PT dinucleotide levels in Donor #5. This source data is associated with **Figure 2E**.

**Table S5.** PT consensus sequence metagenomics.

**Fig. S1.**
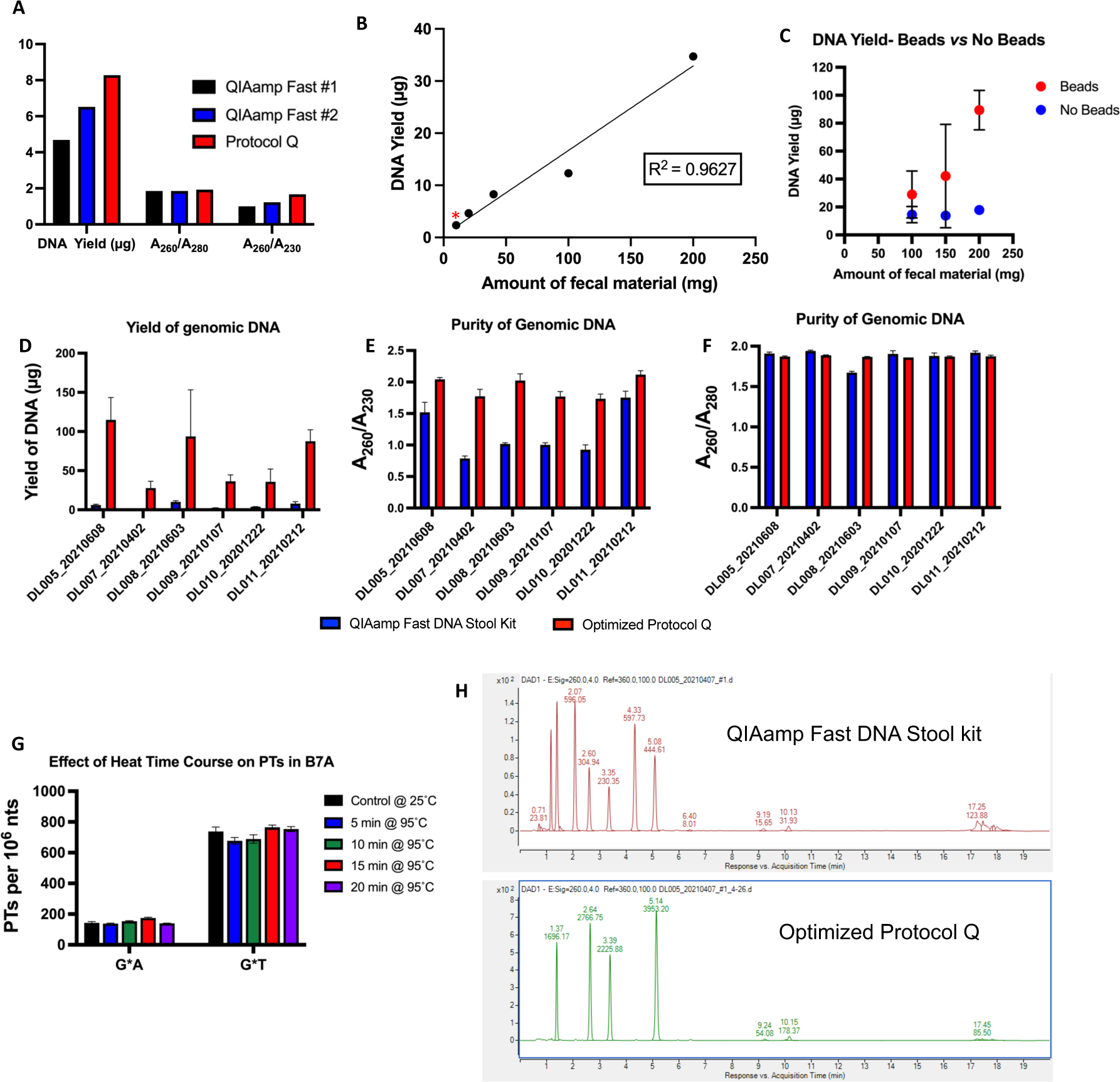
Optimization of Protocol Q for fecal DNA extraction. (**A**) Comparisons of DNA yield and purity for the original, unoptimized Protocol Q (red) and two independent repetitions of the QIAamp Fast DNA Stool Kit (black and blue) with the same human fecal sample. (**B**) DNA yield relative to increasing amounts of fecal material in the same 200 μL volume. The red asterisk denotes the original conditions from Protocol Q, in which only 5% of 20-fold diluted fecal matter was used for DNA purification. The revised method skips the dilution step and uses the entire fecal sample for DNA purification. The yield of DNA from 200 mg of feces thus increased from 2.4 μg to >30 μg. (**C**) DNA yield from human feces when using glass beads (red) or no glass beads (blue) in the revised Protocol Q. (**D, E, F**) Comparison of DNA yield (**D**), A_260_/A_230_ ratio (**E**), and A_260_/A_280_ ratio (**F**) with the QIAamp Fast DNA Stool kit (blue) and the optimized Protocol Q (red). DNA was extracted from human fecal samples from 6 donors using the two different protocols. (**G**) Effect of heat on the stability of phosphorothioates in PT-bearing genomic DNA isolated from *E. coli* B7A and spiked into human feces. PT dinucleotides were analyzed in nuclease-digested fecal DNA analyzed by LC-MS. (**H**) HPLC elution profile (UV detection) of nuclease digests of human fecal DNA isolated using the QIAamp Fast DNA Stool kit (upper) and the optimized Protocol Q (lower).

**Fig. S2.**
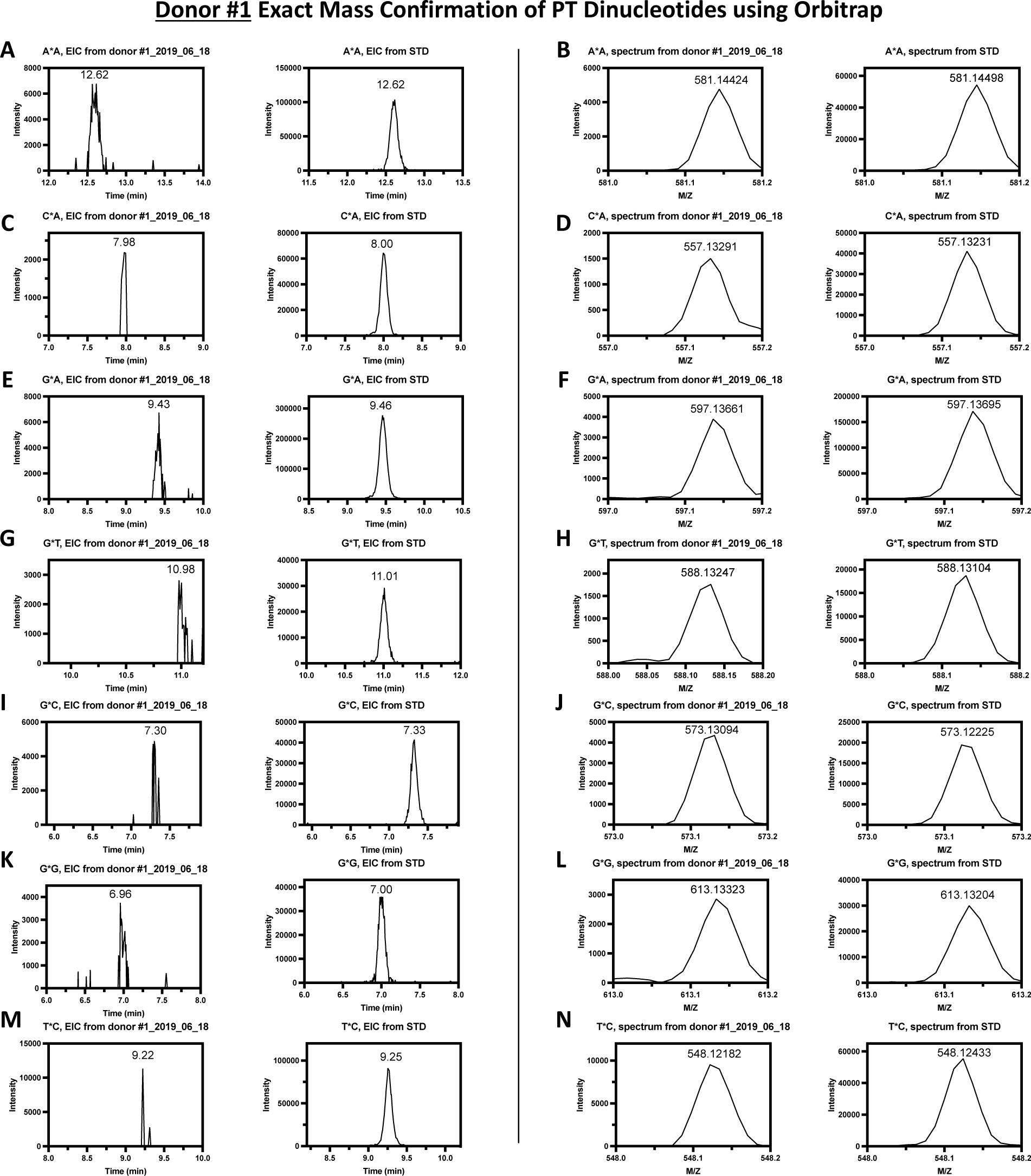
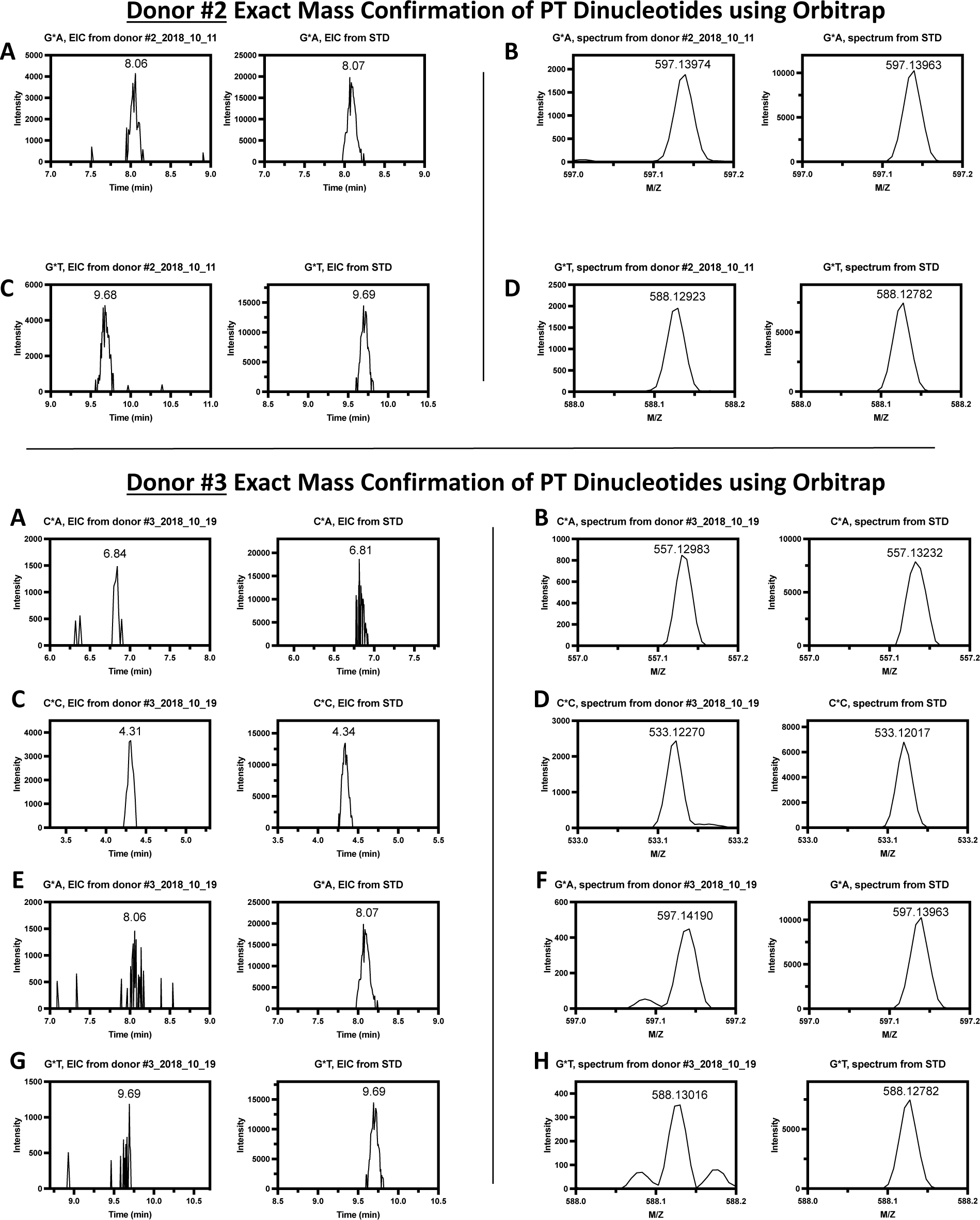
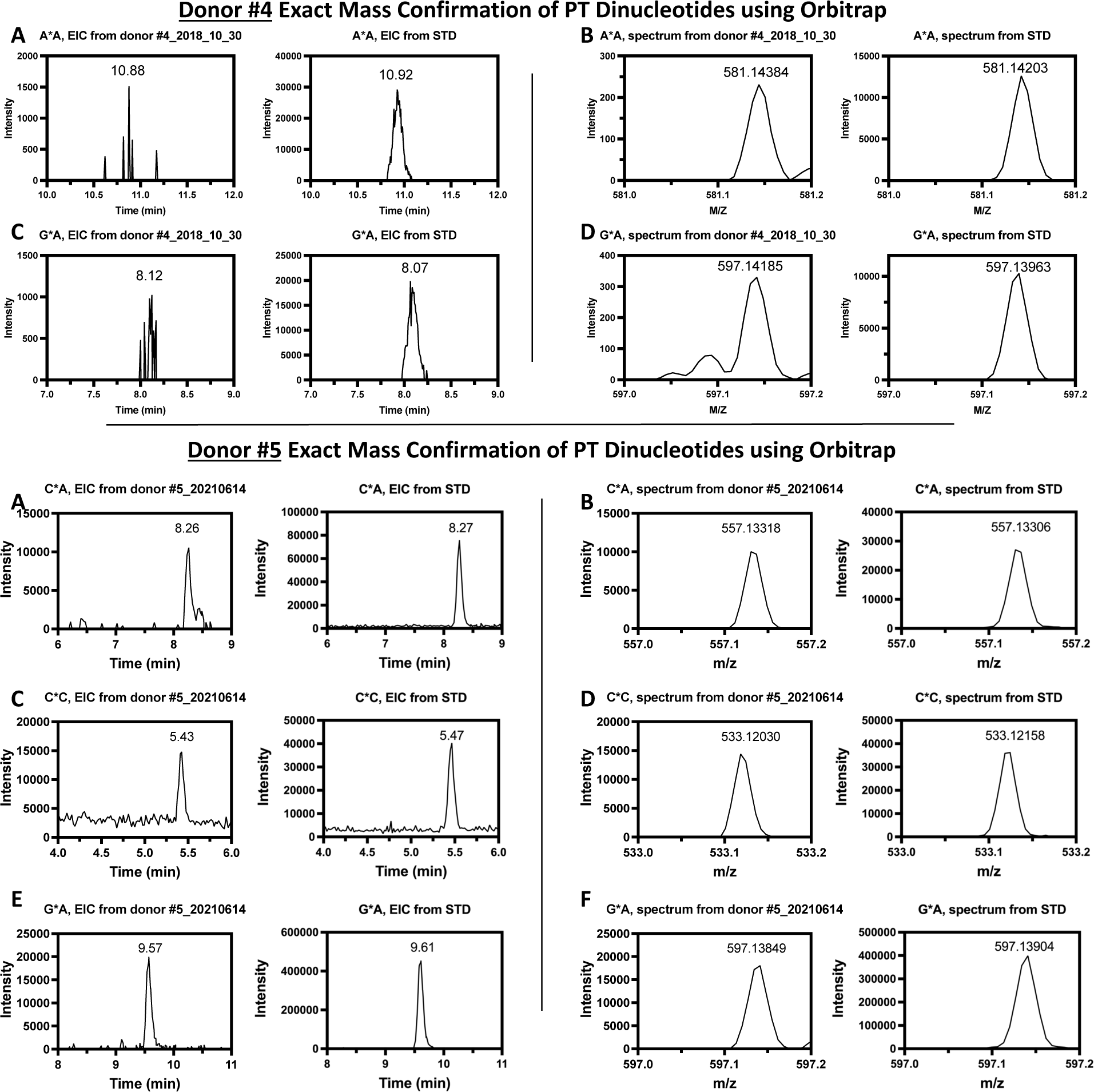
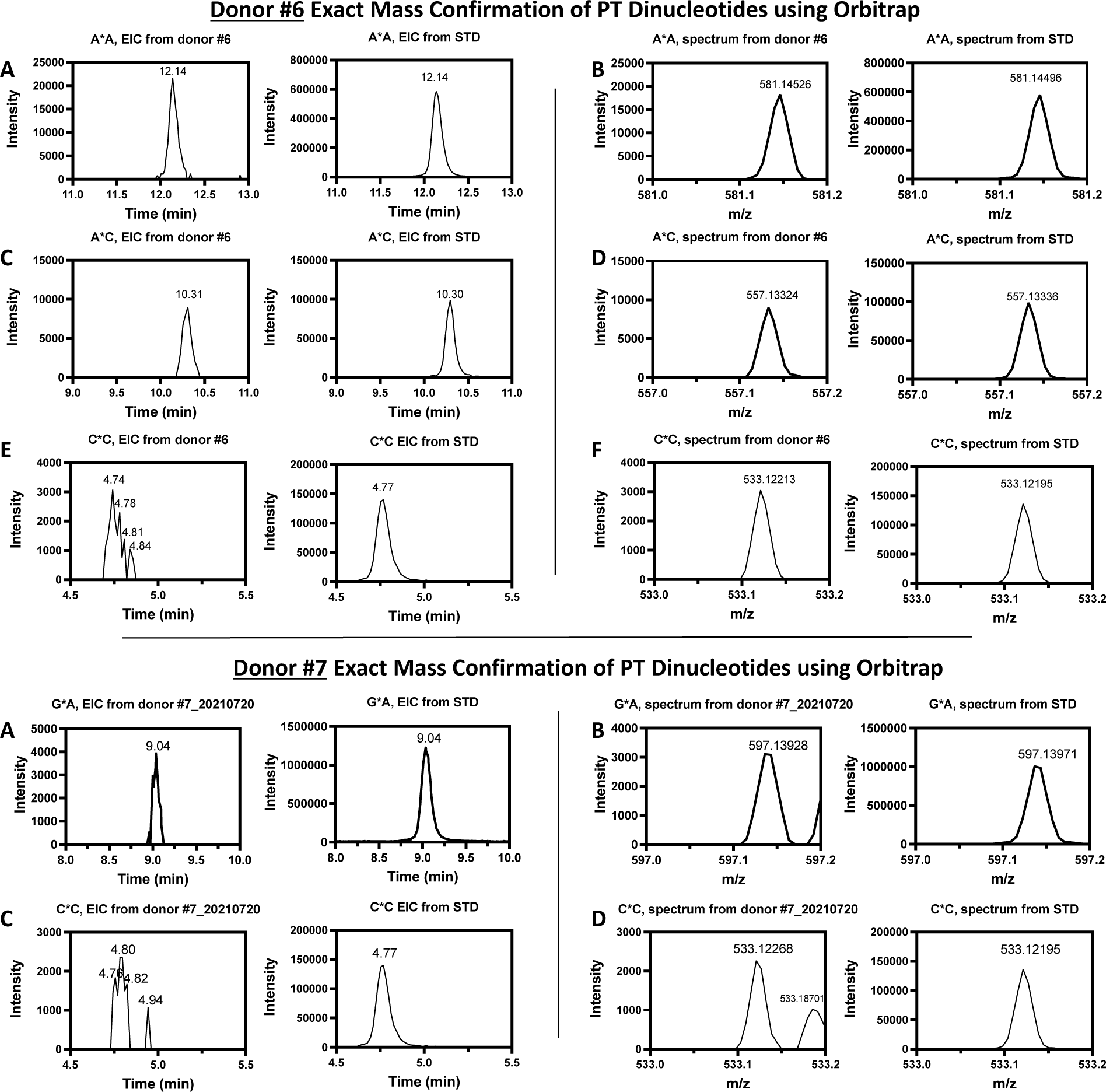
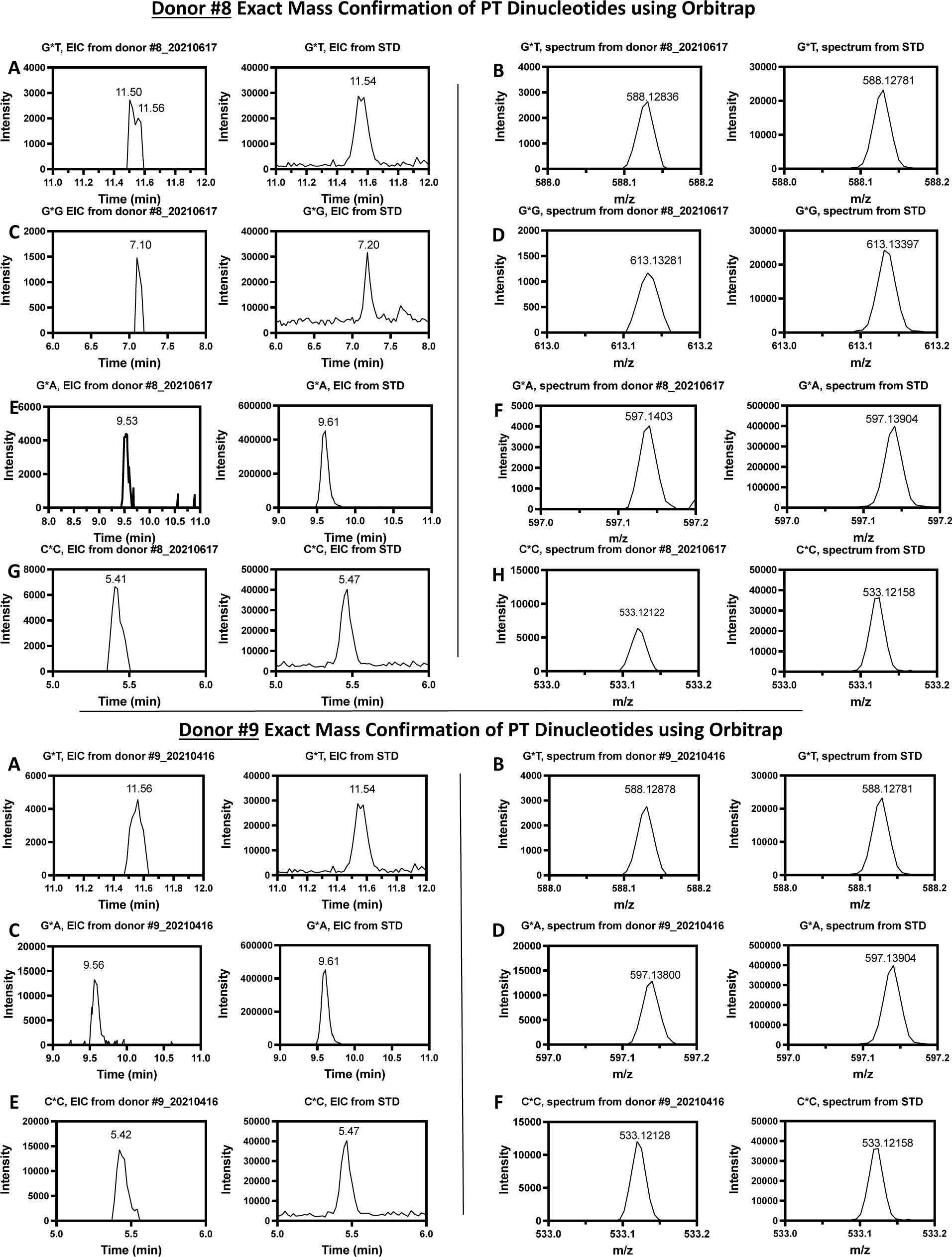
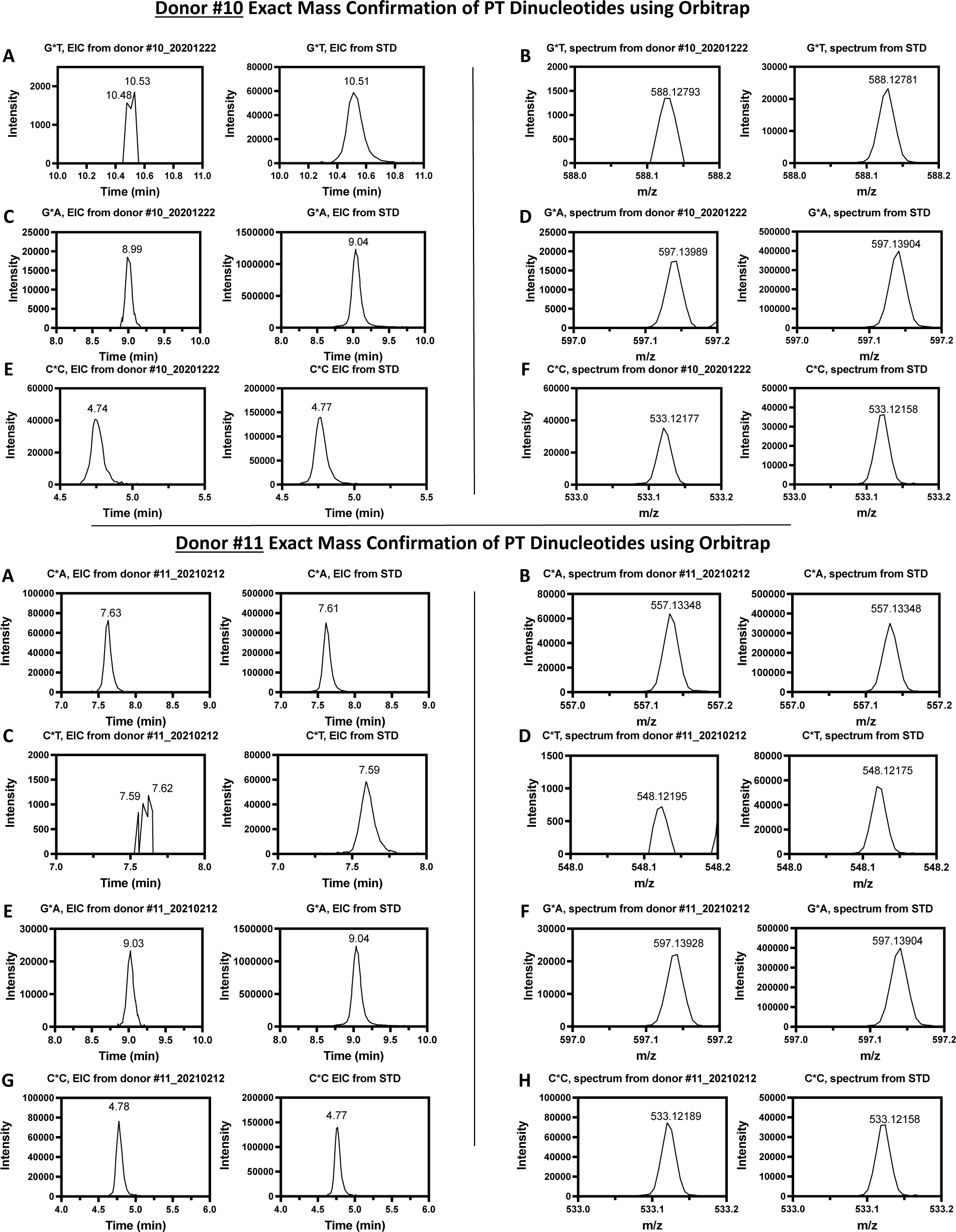
Exact mass confirmation of PT dinucleotides using Orbitrap for donors #1-#11. For each donor, from the left, the first and second columns show extracted ion chromatograms for the putative PT dinucleotide and the relevant standard, respectively, and the third and fourth columns show the mass spectra of the putative PT dinucleotide and the relevant standard, respectively, within 10 ppm.

**Fig. S3.**
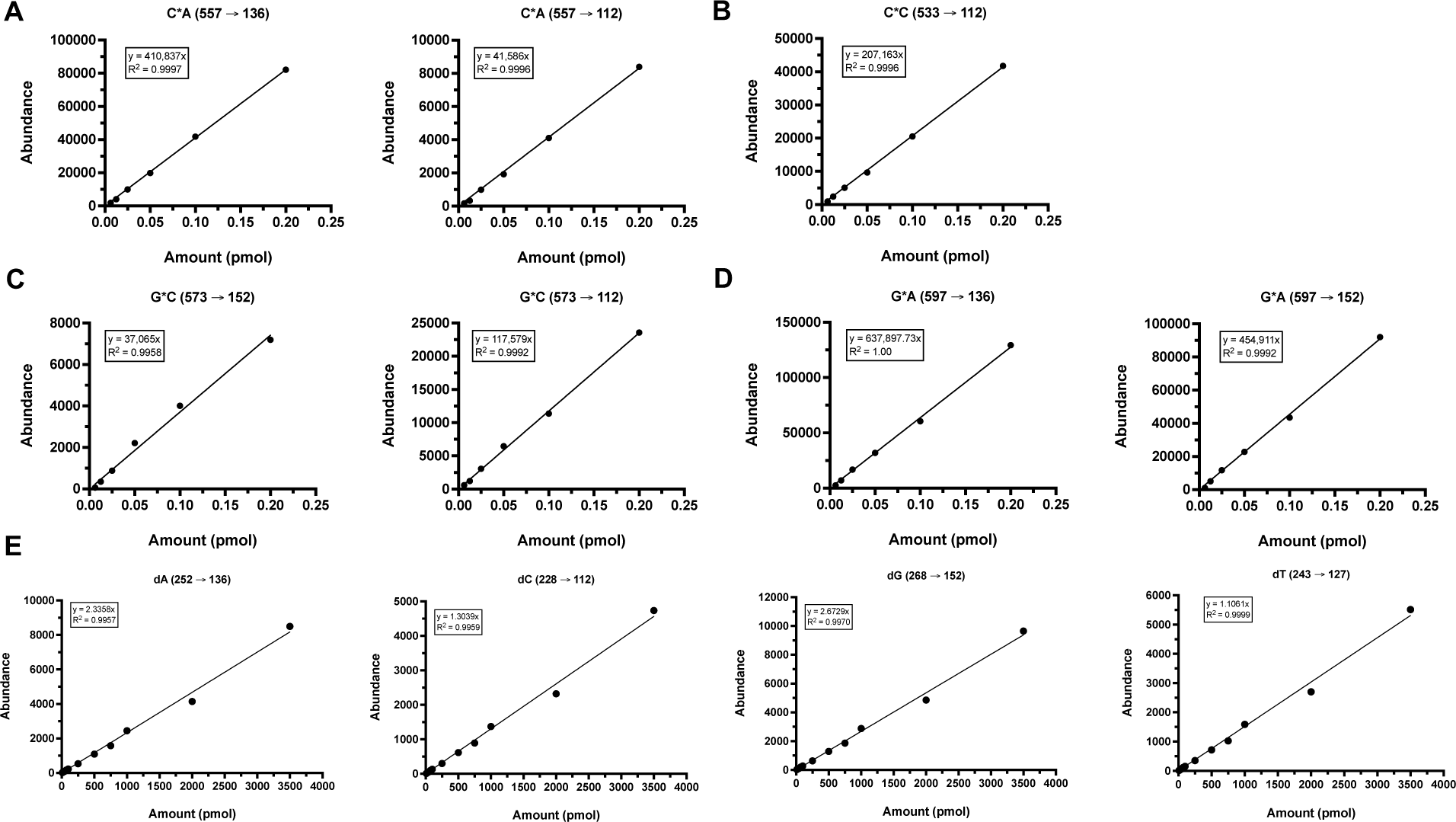
LC-MS calibration curves for PT dinucleotides for donor #5. Calibration curves for the PT dinucleotides C*A. **(A)**, C*C **(B)**, G*C **(C)** and G*A **(D)** were generated by plotting the amount of each PT dinucleotide synthetic standard that was injected onto the LC-MS against the measured abundance for the indicated MRM transition by LC-MS. **(E)** Calibration curves for the 4 canonical deoxyribonucleosides were generated by plotting the amount of each deoxyribonucleoside synthetic standard that was injected onto the LC-MS against the measured UV signal at 260 nm using the diode array detector on the LC-MS. A line of best fit was constructed for each calibration curve, along with the equation describing the line of best fit and the coefficient of determination (R^2^).

**Fig. S4.**
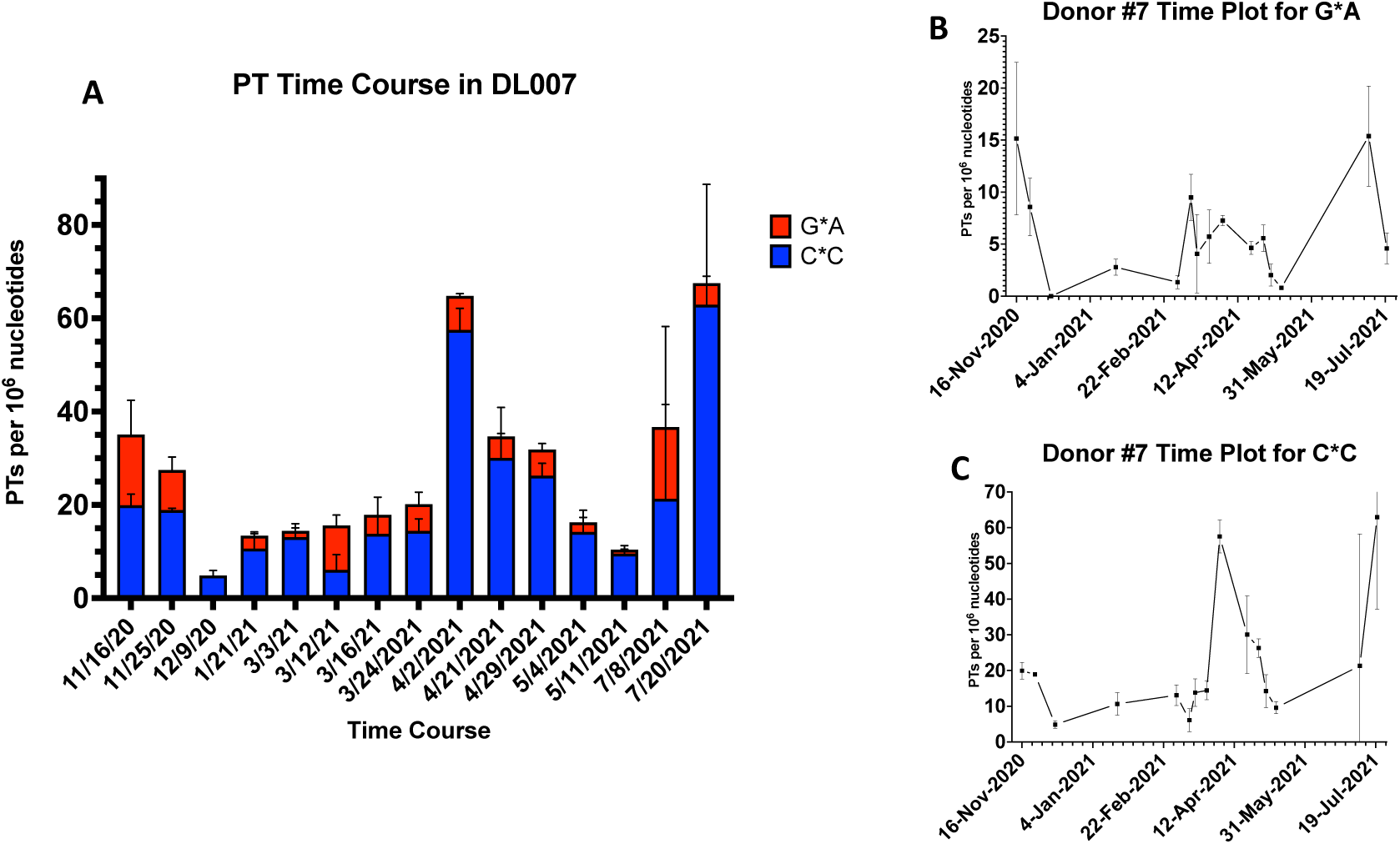
PT dinucleotide time course for donor #7 for. **(A)** total PTs, **(B)** G*A, and **(C) C*C**. Data represent mean ± SD for N=3.

**Fig. S5.**
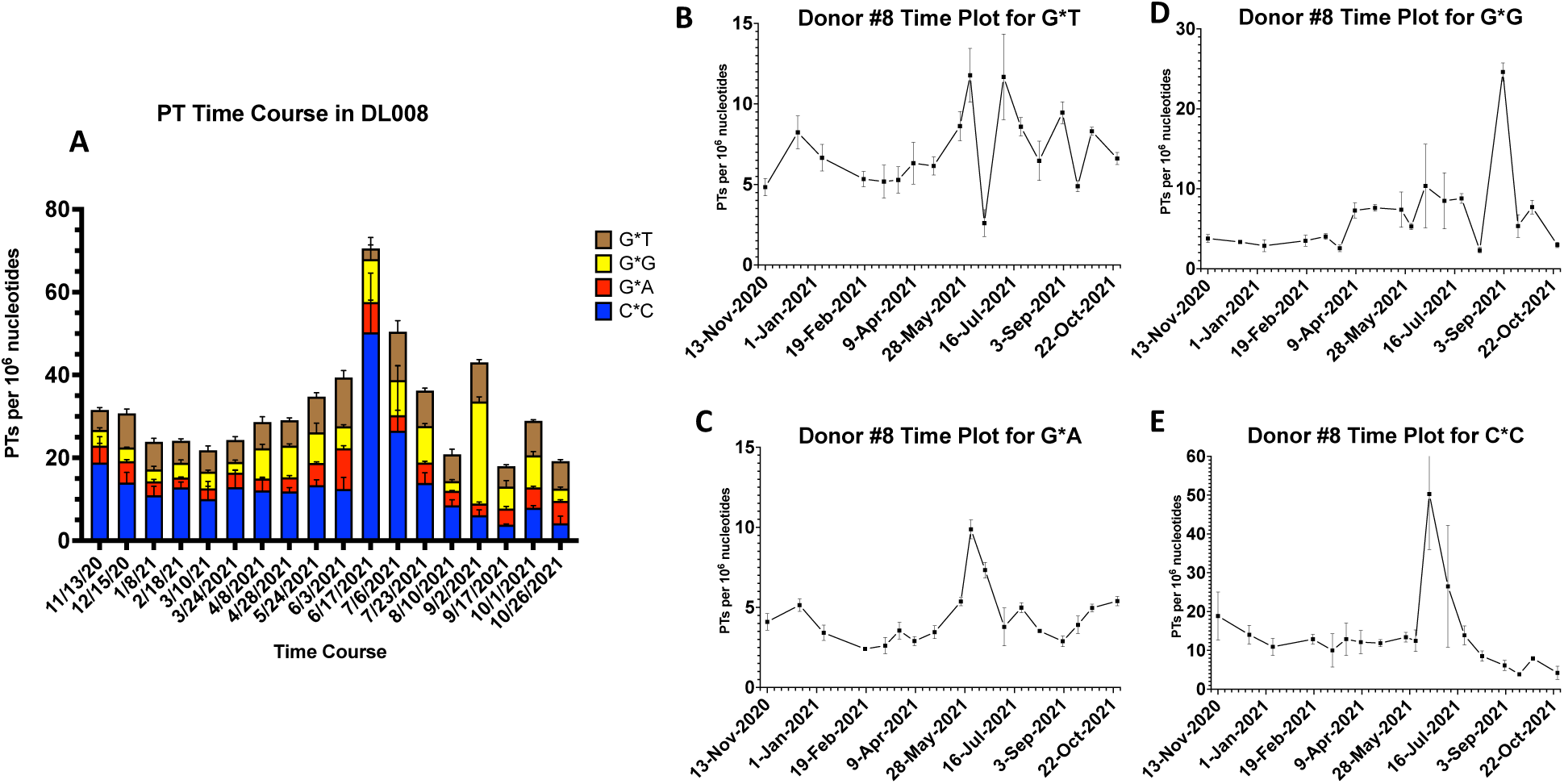
PT dinucleotide time course for donor #8 for. (**A**) total PTs, (**B**) G*T, (**C**) G*G, (**D**) G*A, and (**E**) C*C. Data represent mean ± SD for N=3.

**Fig. S6.**
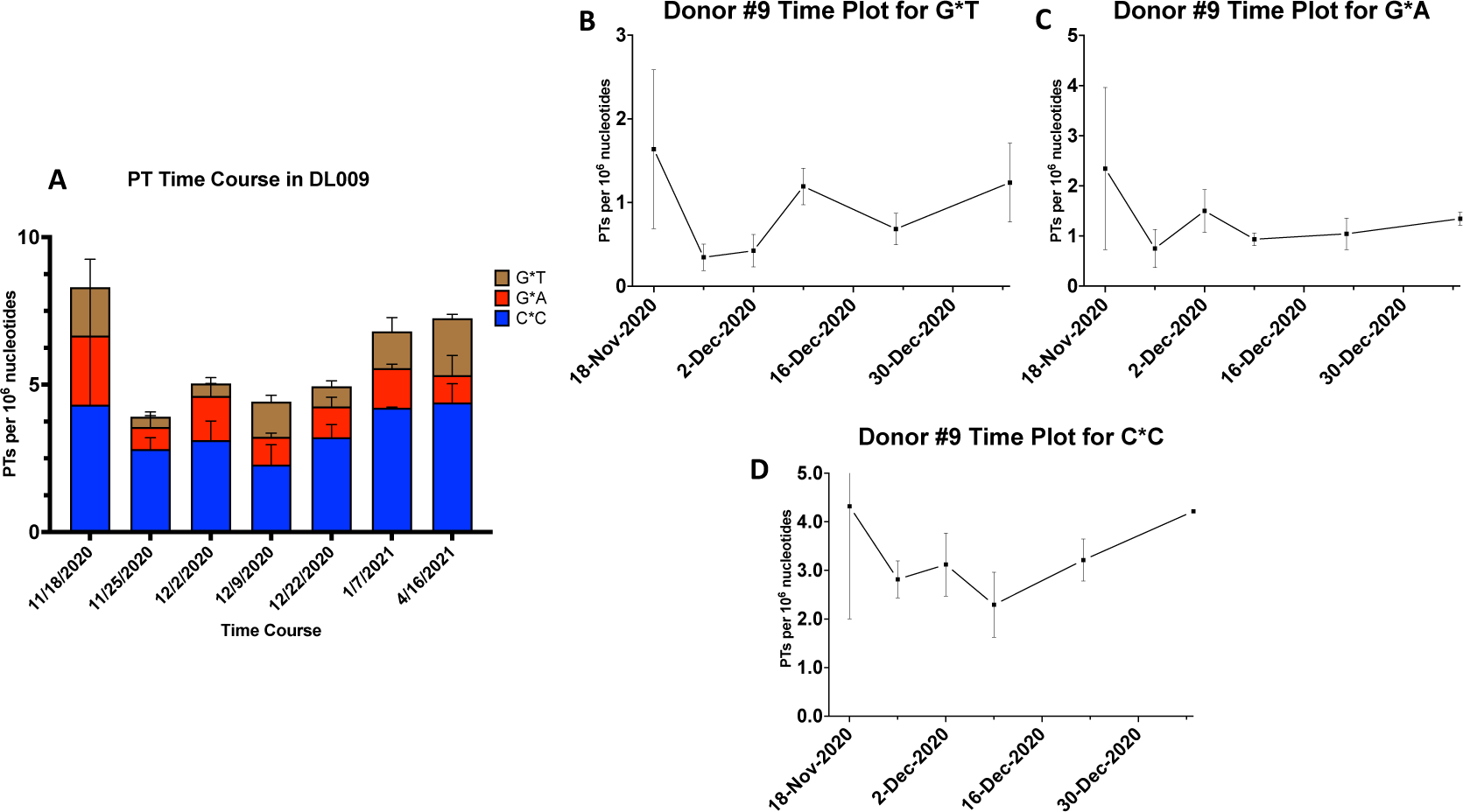
PT dinucleotide time course for donor #9 for. (**A**) total PTs, (**B**) G*T, (**C**) G*A, and (**D**) C*C. Data represent mean ± SD for N=3.

**Fig. S7.**
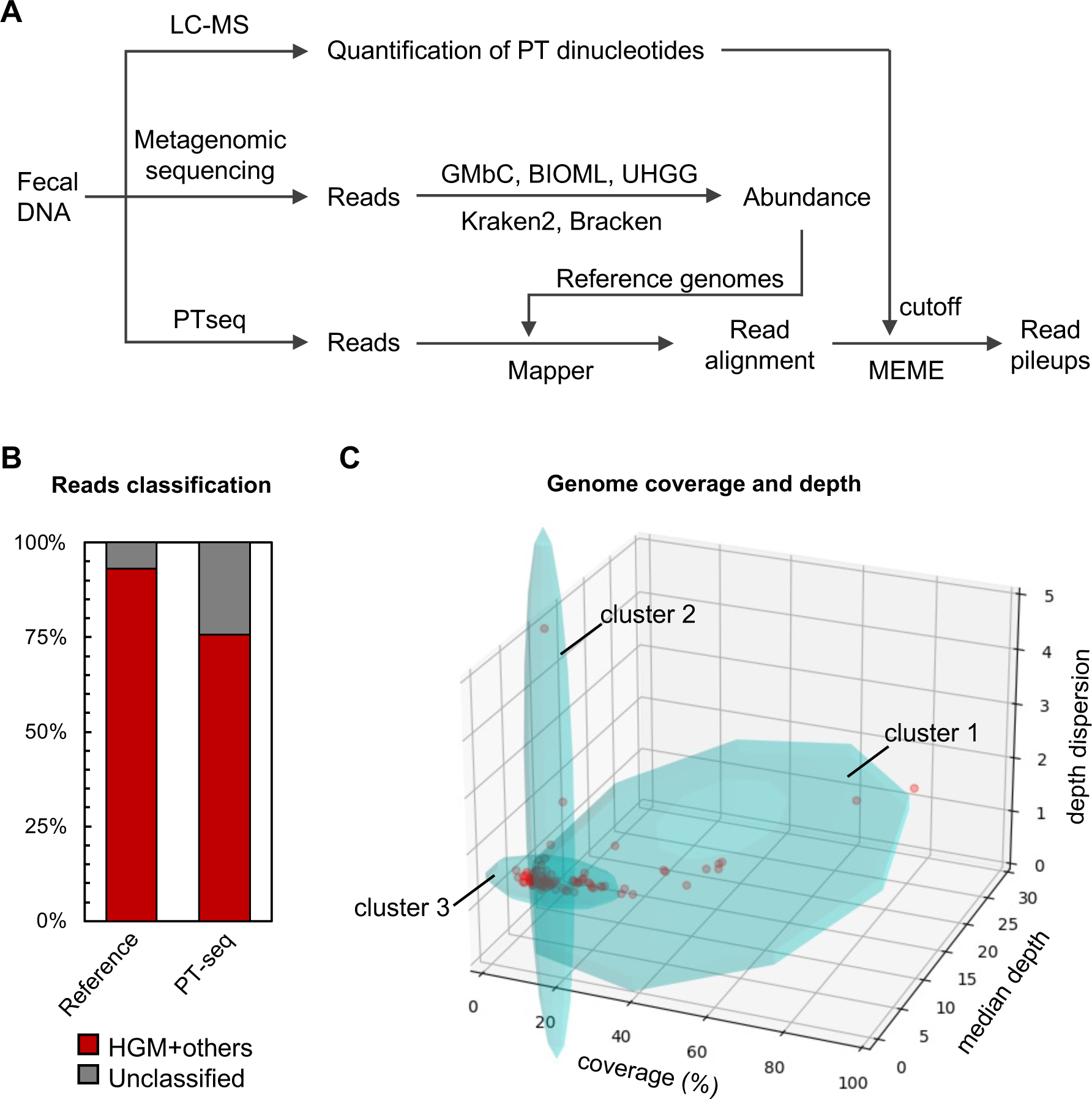
Processing of metagenomic sequencing and PT-seq data. (A) The scheme of investigation of PT landscape of human gut microbiome. Metagenomic sequencing reads were assigned to HGM genomes. PT-seq reads were mapped to the most abundant 100 genomes using Mapper v1.1-beta04. The genomes with above 15% coverage were subjected to MEME consensus motif detection and pileup site analyses, in which the cutoff of read pileup depth was determined by the quatification of PT dinucleotides using LC-MS. (B) The read classification of metagenomic sequencing (reference) and PTseq of human gut microbiome genomes of donor #5. (C) The clustering of the most abundant 100 genomes for PTseq. PTseq reads were mapped to 100 genomes with the most read assigned by Bracken using Mapper v1.1-beta04 (github.com/mathjeff/Mapper). The coverage, median depth and dispersion of depth were plotted and clustered using GMMs. Genomes in cluster 1 were subjected to pileup site analyses.

**Fig. S8.**
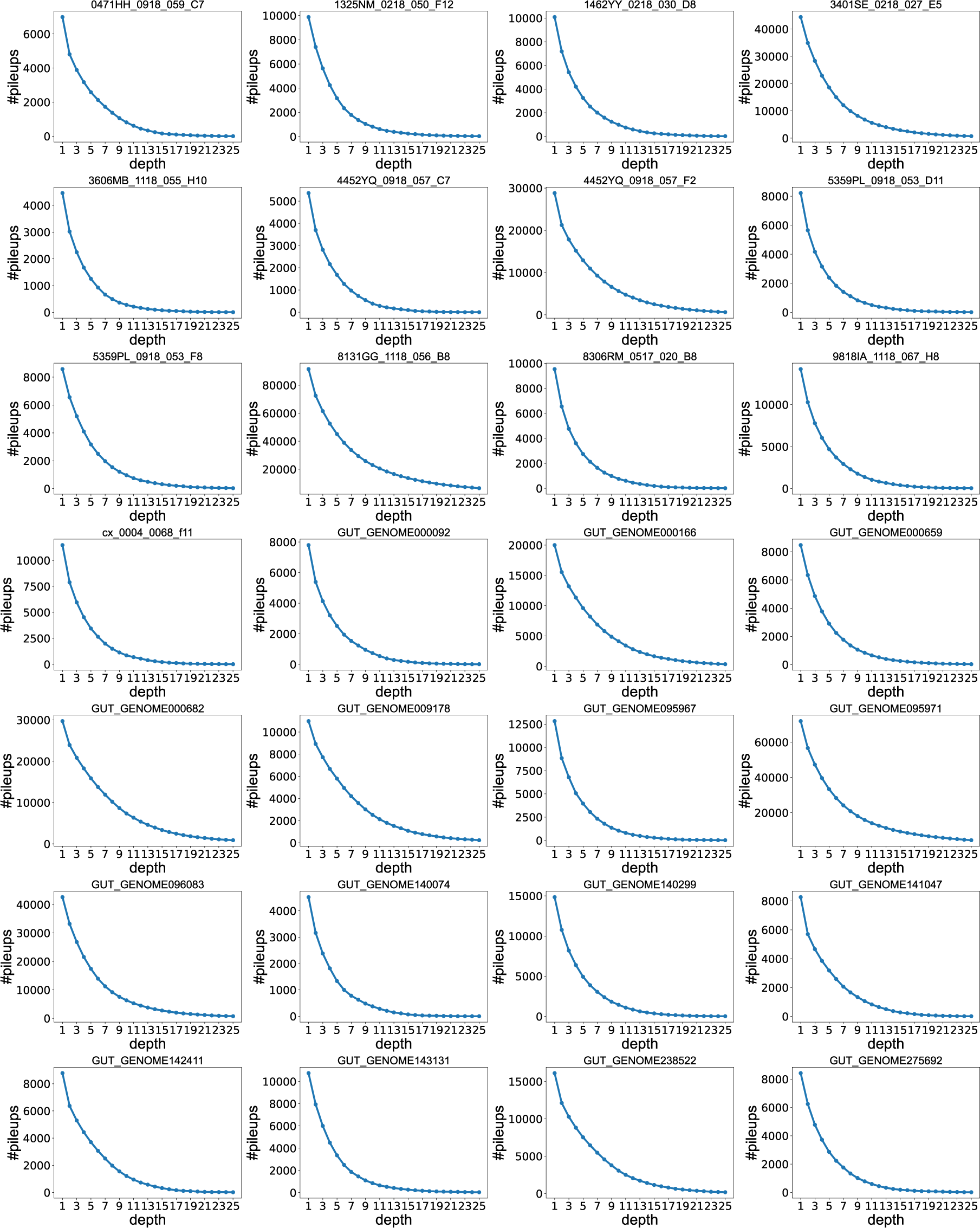
The number of read pileups (y-axis) with an increasing depth cutoff (x-axis) in 28 genomes with >15% coverage. Each graph depicts 1 of the 28 genomes. The number of read pileups decreases with increasing depth cutoff in a negative log-like manner. Reaching a cutoff of 15 converged on 26,507 C*AG, C*CA, C*CGG, G*ATC and G*AGC sites that correlated with the LC-MS levels of three PT dinucleotides (C*A, C*C, and G*A).

**Fig. S9.**
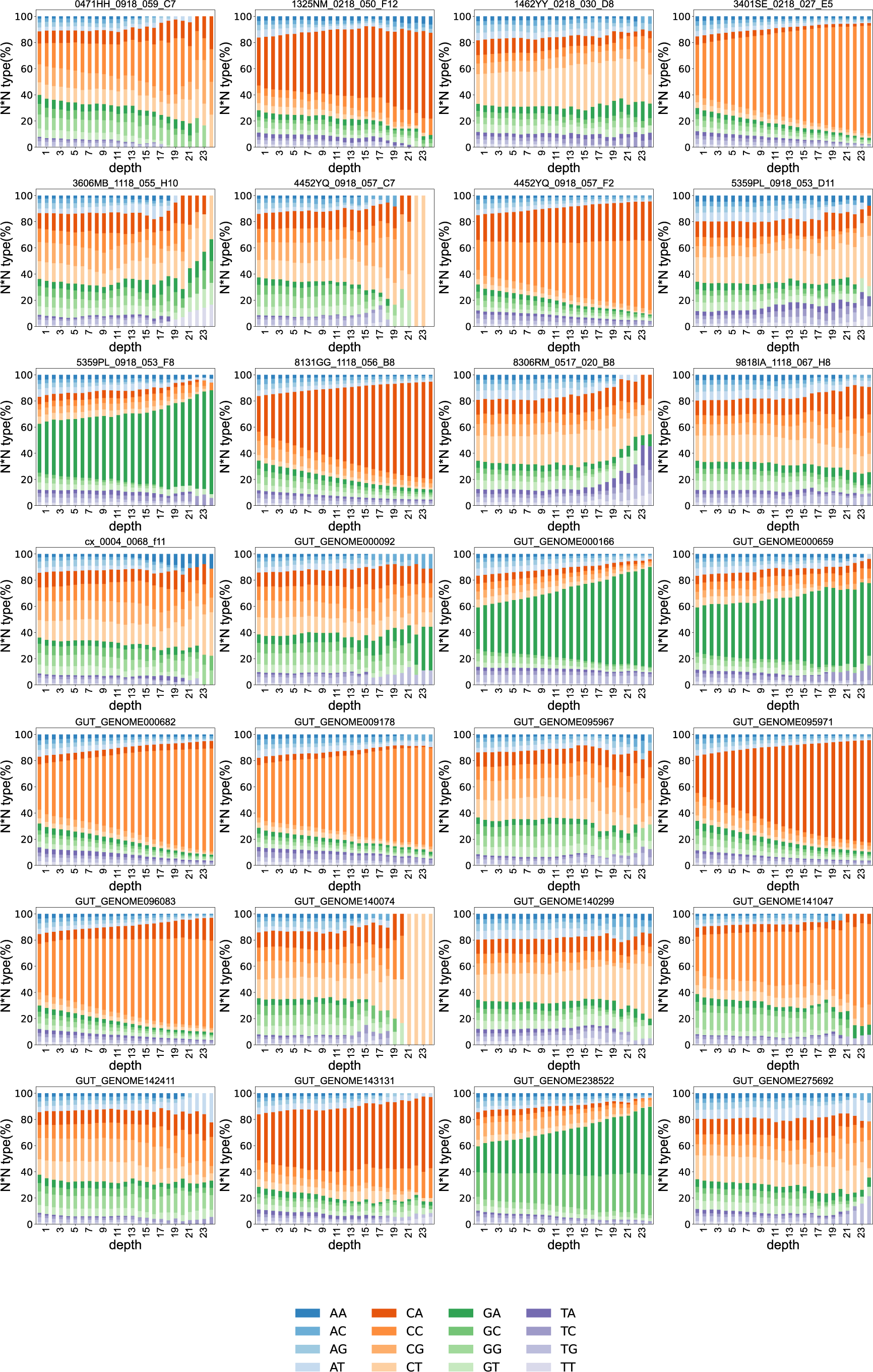
The percentage of read pileups at each of 16 possible dinucleotide sites. The percentage (y-axis) as a function of read pileup depth cutoff (x-axis) in 28 genomes with >15% coverage. Each graph depicts 1 of the 28 genomes.

